# Aging Leads to Altered Physiological Reactivity in Response to Repeated Social Separation Stress in a Nonhuman Primate Model

**DOI:** 10.1101/2025.10.03.680353

**Authors:** Aaryn Mustoe, Jessica Greig, Addaline Alvarez, Clarissa Hinojosa, Jessica Duran, Alanna Melchor, Eden Comer, Juan-Pablo Arroyo, Hillary F. Huber, Donna Layne-Colon, Ektoras Lambrou, Jessica Callery, Luis Giavedoni, Kimberley A. Phillips, Emily S. Rothwell, Adam B. Salmon, Corinna N. Ross

**Affiliations:** Southwest National Primate Research Center, Texas Biomedical Research Institute, San Antonio, TX, USA; Department of Biology, Trinity University, San Antonio, TX, USA; Department of Psychology, Trinity University, San Antonio, TX, USA; Department of Neurobiology, University of Pittsburgh School of Medicine, Pittsburgh, PA, USA; Barshop Institute for Longevity and Aging Studies and Department of Molecular Medicine, University of Texas Health San Antonio, San Antonio, TX, USA; Geriatric Research Education and Clinical Center, South Texas Veterans Healthcare Center, San Antonio, TX, USA

**Author notes:** Corresponding Author: Aaryn Mustoe, Southwest National Primate Research Center, Texas Biomedical Research Institute.

**Keywords:** social aging, stress reactivity, cortisol, inflammation, metabolism, marmosets

## Abstract

Social relationships are critical for maintaining physical health and psychological wellbeing. Since nearly 1 in 4 adults aged 65 years or older are socially isolated, there is a strong need to understand how repeated social stress negatively impacts health outcomes. Using a nonhuman primate model of social aging (i.e., marmosets), we examined whether individuals transitioning into old age (“peri-geri”) or individuals who were already geriatric (“very-geri”) showed differences in measures of hypothalamic-pituitary axis (HPA) activity, markers of metabolic and immune function, and reunion social behavior in response to repeated social separation challenges (SSC). We found female marmosets, especially peri-geri females, had higher HPA reactivity and better HPA recovery than male marmosets, but this difference diminished in older, very-geri marmosets. HPA activity was correlated with multiple outcomes including locomotive behavior and grooming, changes in blood glucose levels, and neutrophil counts. Moreover, marmosets who approached their partner more and were groomed less during reunions had higher cortisol levels the following day. Interestingly, we found two distinct HPA profiles among our marmosets with half showing strong HPA responses (reactors) and the other half showing little or no HPA response (non-reactors). Non-reactors had less weight gain/more weight loss; elevated levels of calcium, phosphorus, and white blood cells; and received less grooming and social contact time during reunion. Overall, old-aged marmosets who show attenuated HPA responses may have different vulnerabilities to negative health and behavioral outcomes during social stress, and male marmosets appear more likely to present with this HPA non-reactor phenotype earlier in aging.

## INTRODUCTION

Social relationships are critical for maintaining physical health and psychological wellbeing throughout the lifespan. Many reports examining the association between social function and physical health have indicated that the presence of high social connectiveness and strong social relationships are associated with at least a 50% reduction in mortality rates [1], an effect size that is greater or comparable to other well-established lifestyle factors associated with mortality including cessation of smoking, reduced alcohol consumption, and increased physical activity; Moreover the accumulation of stressors associated with the disruption of social relationships has been shown to negatively impact immune function, increase inflammation, and accelerate aging processes [2–3] leading to increased risk of disease [4–5]. Fundamental to this link between social function and overall health is the fact that disruption of social relationships and social functioning induce a chronic state of physiological and behavioral stress. Characterizing biobehavioral reactions to social isolation stress in aging is crucial for development of preventative and intervention strategies to maintain positive health and wellbeing outcomes.

Across the lifespan, aging introduces additional challenges in maintaining positive social function due to age-related decline in abilities such as mobility, vision and hearing, and cognition, each of which can make it more difficult for maintaining social relationships and make individuals more vulnerable to stress and disease. Recent reports have indicated that about 1 in 4 adults aged 65 and older are socially isolated [6], and research and meta-analyses clearly demonstrate an association between poorer quality social relationships and increased levels of chronic inflammation [7–8]. This is a significant global concern as the Global Burden of Diseases, Injuries, and Risk Factors Study (GBD) has shown that over 50% of all deaths are attributed to some form of inflammation [9]. When considering concentrations of a wide range of inflammatory biomarkers, the pattern of changes in inflammation responses to prolonged “stress” and inflammation during “aging” are remarkably similar [10].

Activation of physiological and behavioral responses via the hypothalamic-pituitary-adrenal (HPA) axis is essential for survival and resilience during events or periods of stress. However, chronic exposure to stress hormones can predispose to psychological, metabolic and immune alterations, and this may be especially important in older populations where changes in cognitive, metabolic, and immune function are magnified. Aging has been associated with decreased sensitivity of glucocorticoid negative feedback in the HPA axis and altered circadian rhythm cyclicity in both nonhuman primates (NHPs) and humans [11–12]. Specifically aging has been linked to increases in evening levels of circulating glucocorticoids and basal glucocorticoid levels overall, but evidence is mixed as to whether aging is associated with higher or lower HPA reactivity [13–14], and in many cases, these findings are sex-dependent with females typically showing greater HPA sensitivity [15–16]. However, it is unclear what factors may contribute to age-related changes in HPA function, including the form or type of stressor(s) (social vs. nonsocial), the magnitude and duration of stressor(s) (acute vs. chronic), and the overall metabolic, cognitive, and immune health status of the participants, all of which represent a major gap in our understanding of how social disruption affects HPA activity (and vice versa) during aging.

Common marmosets (*Callithrix jacchus*) are an emerging and important primate model to examine physiological and behavioral mechanisms that underlie age-related changes. Marmosets are small-bodied arboreal primates found across forest and forest-edge habitats in Brazil, ranging from diverse environments such as the Atlantic coastal forests and the xeric shrublands of the Caatinga region. The social lives of marmosets consist of small social groups usually containing a single dominant breeding pair that are similar to extended family groups in humans [17–18]. One of the hallmarks of human social relationships is the bond between romantic partners, i.e., a “pair-bond” [19]. Marmosets are one of the a few nonhuman primates that show many attributes that are highly similar to human social relationships [20–21]. Marmosets display characteristics of a “pair-bond” which includes preferential affiliation toward their partner, coordination of social behavior to each other, increased physiological and behavioral stress during social separation, and increased mate-guarding in the presence of other same-sex intruders [22–26]. Given that marmosets are highly sensitive to social stress in the context of their social relationships with others [27–28], marmosets offer significant potential advantages as an NHP model to determine biobehavioral mechanisms associated with social stress and aging.

In this study we were interested in examining whether aging and geriatric marmosets have distinct physiological and behavioral changes as a result of repeated social separations that are distinct from what we know previously in adult and younger aged populations. The primary design of this project used a social separation challenge (SSC) paradigm, which functions as a period of repeated, short-term social separations that includes an individual being removed from their homecage and being isolated in a room away from their social partner without any visual access to other conspecifics. The goal of the SSC is to mimic the repeated, prolonged, and potentially negative components of stress by exposing individuals to the accumulation of social stressors as individuals transition from adulthood into old-age [29–30]. For example, geriatric humans may experience an accumulation of social isolation due to hospital stays, deaths of family or friends, living alone, and/or leaving the home with decreasing frequency. Individual marmosets in this study fell into one of two categories of aging. One group, “*<u>peri-geri</u>*”, included marmosets aged between 6 to 9-years-old, an age considered to be the timeframe when individual marmosets are transitioning from middle into geriatric age. The second group, “*<u>very-geri</u>*” are marmosets aged 9+ years-old, and are considered geriatric, which is an age when many of the changes in phenotypes of aging, including behavior, typically have emerged [31–32]. Overall, our goal was to examine whether aging marmosets show differences in physiological and behavioral responses during repeated SSCs, and whether other behavioral or physiological factors including markers of hypothalamic-pituitary axis (HPA) activity, immune function, and social behavior during social reunion, would enhance or mitigate measures of stress reactivity or stress recovery during repeated SSCs.

## METHODS

### Subjects

Fourteen common marmosets (*Callithrix jacchus*) participated in this study. All animals were socially housed as continuous full-contact female-male pairs (cohabitation length mean ± SD = 3.15 ± 2.78 yrs; cohabitation length range = 0.43 – 8.78 yrs), at the Southwest National Primate Research Center (SNPRC), Texas Biomedical Research Institute, an AAALAC accredited institution. These 14 marmosets were composed of 8 females and 6 males. Animals were housed in male-female pairs, and only one individual per pair (n = 14) was removed from the homecage for the SSCs in this study, meaning that each of the 14 individuals tested during SSCs came from a unique male-female pairs. Of note, cohabitation length or number of previous partners was not significantly associated with any physiological or behavioral data outcomes. Marmosets were split into two discrete age categories based on the principle that there are potential important categorical differences in physiology and behavior between animals who are transitioning from adulthood into old age (*peri-geri*) and animals that are already considered to be “old age” (*very-geri*). These age groups resulted in an age range as follows (range; mean ± standard deviation): *Peri-geri*: 5.96-8.89; 7.54 ± 1.07 yrs and *Very-geri:* 9.32-13.11; 10.92 ± 1.38 yrs. A large multi-institution study calculated median lifespan of a common marmoset (n=831) in captivity across primate research centers as 5.97 (5.41–6.74) for males and 5.31 (4.92–5.66) for females with a maximum lifespan reported at ∼19 years at SNPRC [33]; these median life expectancies are thought to align with healthspan due to the common practice of humanely euthanizing research animals that have developed chronic diseases affecting quality of life. The general consensus is around the age of ∼8 years old, marmosets typically begin to show signs of physiological age-related decline [31]. While old-aged animals are more likely to have underlying health conditions such as metabolic or immunological differences compared to younger adults, all animals enrolled in the study were healthy without any explicit clinical symptomology (*i.e., ongoing weight loss, anemia, gastrointestinal distress, behavioral lethargy, etc.*) that would differ from otherwise healthy adults. Males and females were evenly split across these two age groups with 3 males and 4 females in each age group respectively. Animals were maintained under standardized marmoset husbandry conditions [34], and received base diets (Harlan Teklad marmoset purified diet and Mazuri marmoset diet) as previously described [35]. The study was approved by the Texas Biomedical Research Institute Animal Care and Use Committee, and adhered to the American Society of Primatologists (ASP) Principles for the Ethical Treatment of Non-Human Primates.

### Social Separation Challenge (SSC) Procedure

All 14 marmosets underwent ten individual SSC procedures over a period of five weeks, with two SSCs over a given week, with always at least two full days between individual SSC sessions. A schematic of the SSC is shown in **Figure 1**. Each SSC session began at approximately 0700AM, with an initial urine collection that served as their first-void baseline. At approximately 0730AM, the individual marmoset would be transported from their homecage room to a separate room where the animal would be physically isolated from their long-term social partner, without visual access to any other marmoset. The isolation enclosure included free access to water, food, a wooden branch, and a transparent bucket affixed inside their enclosure. Immediately after the individual as transported to a separate room at 0730AM, a video camera was turned on to record the isolated marmoset for ∼30 minutes, which constitutes the “isolation behavior”. At the top of each hour from 0800AM to 1100AM, a trained researcher would quietly enter the room to noninvasively collect urine that accumulated on a plastic bedding sheet that was placed beneath their enclosure. These enclosures are smaller versions of standardized marmoset husbandry previously described [34]. At 1100AM, the isolated marmoset would be transported back to their homecage and a 20-minute live-scored focal reunion observation was recorded on the isolated individual as they were reunited with their long-term social partner, which constitutes the “reunion behavior”. On the following day at 0700AM, another urine collection was performed to provide a day-after first-void baseline sample. **Figure 1** also shows a timeline and description of the overall SSC procedure. All urine samples were initially frozen at −20°C and transferred for storage at −80°C until assay.

**Figure 1.**
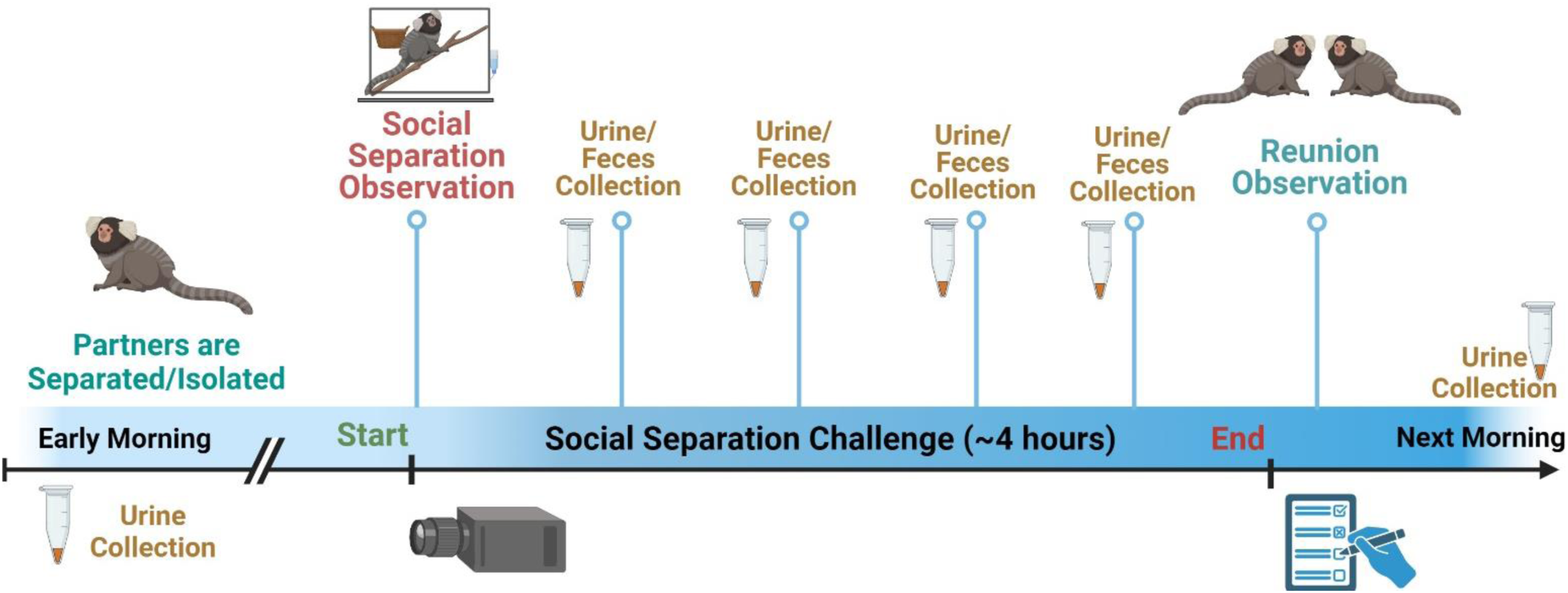
Schematic and timeline of procedures and sample collection for an individual Social Separation Challenge (SSC). SSCs were repeated 10 times for each individual marmoset twice per week for 5 weeks. First or baseline urine samples (both day-of and day-after) were collected at ∼0700AM upon awakening. SSCs lasted from 0730-1100AM with video isolation observations occurring from 0730-0800AM and the live homecage social reunion observations occurring immediately after the SSC (at 1100AM).

### Behavioral Observations

A description and ethogram for both the isolation behavior and the reunion behavior observations are shown in **Table 1**. During the initial social separation observation, *<u>isolation behavior</u>* was scored by trained research staff for the first 5 minutes of the video observation and the last 5 minutes of the video observation. This was done based on the *a priori* assumption that the first 5 minutes of the isolation observation likely included behavioral responses associated with both the separation from their social partner and the physical transportation from one-room to another room, while the last 5 minutes of the video observation likely reflects only behavioral responses associated with the separation from their social partner and would have likely behaviorally recovered from the physical transportation over the first ∼25 minutes of the isolation period. Before performing any data analyses between isolation behavior and HPA activity, we evaluated whether there were behavioral differences between the first 5 and last 5 minutes of the video isolation observations and whether these differed by factors in our study (age and sex). We found that the expression of some isolation behaviors (particularly locomotor behaviors) was higher during the first 5 minutes of isolation, which is consistent with the idea that marmosets were more active/aroused during the first 5 minutes of their isolation compared to the last 5 minutes; Importantly, though, this effect did not vary by sex or age; thus, unless otherwise reported, isolation behavior data was analyzed as averaged across the first and last 5 minutes. *<u>Reunion behaviors</u>* were scored by a live, in-person observer for the full 20-minute observation. Behavioral data is reported as total # or duration of behaviors scored per observation.

**Table 1.**
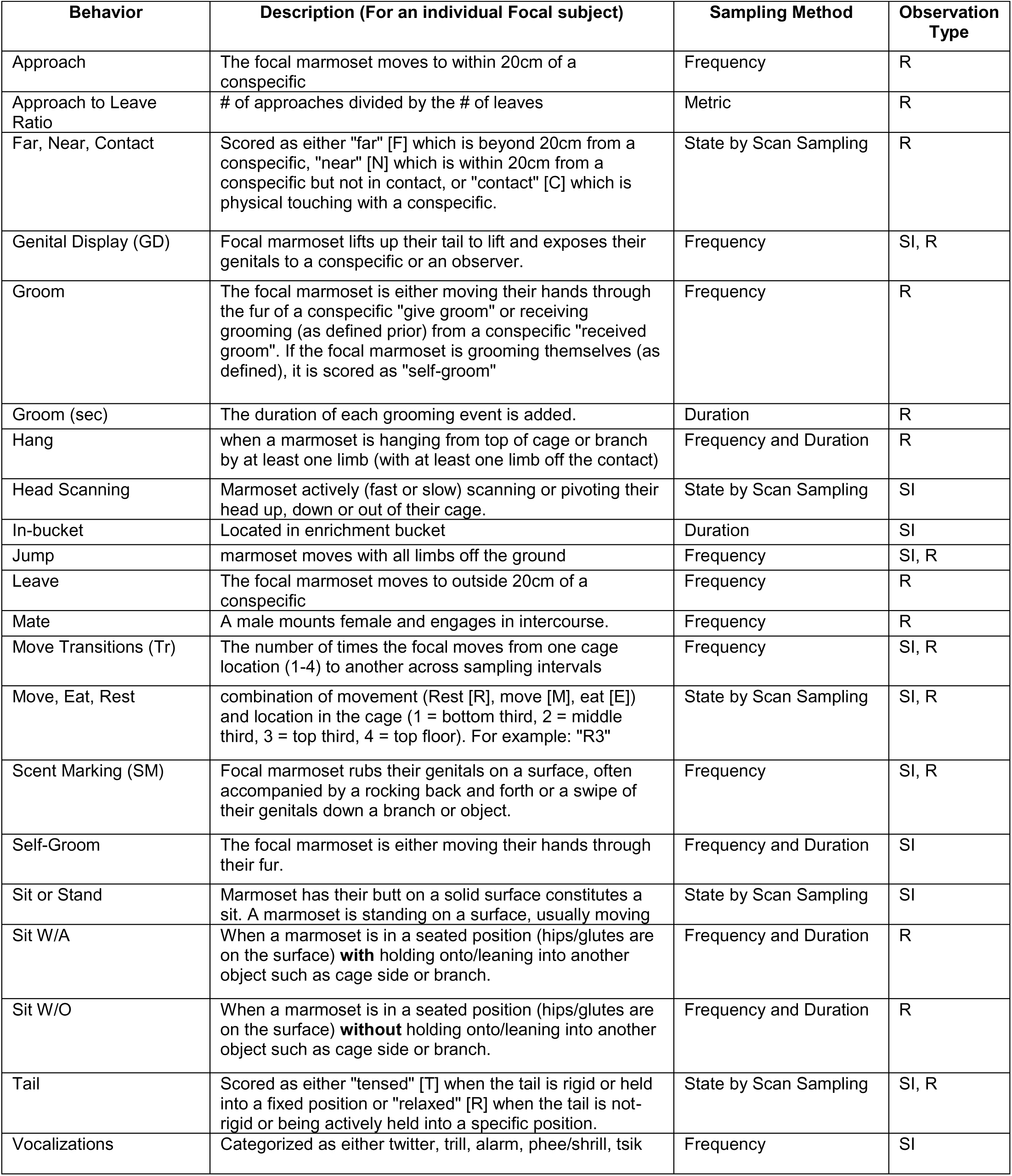
Description of behavioral ethogram for social separation observations (isolation behavior) and reunion observations (reunion behavior). Ethograms used interval sampling for 20 sec periods. State equals presence or absence at each 20s interval. Frequency equals total # of observed behaviors. Duration is the total time spent in behavior in seconds. Observation type: SI = Social isolation observation; R = Reunion observation

### Urinary Cortisol Enzyme Immunoassay (EIA)

We used EIAs to quantify urinary cortisol concentrations. 96-well flat-bottom immuno MaxiSorp microtitre plates were coated with cortisol antibody (diluted to 1:25,000 in bicarbonate coating buffer), and incubated at 4°C for between 12-48 hrs. After the cortisol antibody incubation, 50 µl of PBS was added to each well, followed by 50 µl of the diluted urine samples (1:6400 in distilled water), cortisol standards (cortisol standards were diluted in PBS and ranged from 1000 to 7.8 pg/well) to corresponding wells using a consistent plate template layout, and followed by 50 µl labeled cortisol conjugated with horseradish peroxidase (HRP) (diluted 1:35,000 in PBS) to each well. Stock cortisol (R4866) antisera and conjugated-HRP hormone were originally developed by Coralie Munro, University of California Davis Clinical Endocrinology Laboratory. After the 50 µl of HRP-conjugated cortisol was added, the plates were incubated in a dark chamber at room temperature for ∼2 hrs. Following this incubation, free and bound hormones were separated by automated washing of the plate three times with EIA wash buffer. Following the plate washing, 100 µl of an EIA substrate (ABTS, H_2_O_2_) was added. Absorbance at 405 nm was measured in a microplate reader when an B_0_ optical density of ∼1.0 was reached. Inter-assay CVs for high and low concentration pools were 14.4% and 11.9%, and intra-assay CVs were 7.8% and 9.1%. All samples and standards were run in duplicate. Validation of this assay for marmosets has been performed previously [24]. The mass of cortisol is expressed in µg/mg of creatinine as measured using a standard Jaffé reaction colorimetric assay [36], and is used to account for variable fluid intake/dilution of urinary substrates.

### Cytokine/Chemokine Analyses

We measured plasma at four time points including once 2 weeks prior to the start of the SSCs, once during the middle of the 5-week SSC period, once at the end of the 5-week SSC period, and again 2 weeks after the final SSC was completed. Cytokine concentrations in plasma were assessed for all the subjects by using the Luminex system as validated for marmosets and other nonhuman primates [38–39] (New World Monkey Immunoreagent Resource, https://www.trinity.edu/sites/nwmimmunoreagents). The assay included evaluation of the following 8 analytes: CC chemokine ligand 3 (CCL3), CCL4, granzyme B, interferon gamma (IFN-g), interleukin-6 (IL-6), IL-10, interferon-induced protein 10 (IP-10, CXCL10), and tumor necrosis factor-alpha (TNF-a). Only CCL4 and IP10/CXCL10 were used for data analyses in this study as each of the other analytes were concentrations that were below the assay sensitivity.

### Blood Chemistry Panel and Complete Blood Counts (CBC)

Each marmoset received an annual physical exam as a part of routine colony health monitoring, with one occurring prior to enrollment in this study, and another occurring after their participating in the SSCs. The nearest blood chemistry panel and CBC data before and after the SSC was used for these analyses. For the annual physical exam animals are fasted in the morning (removal of food at ∼0800AM), given a dose of 20 mg/kg of ketamine intra-muscular for sedation, and evaluated by a veterinarian. To evaluate CBC and blood chemistry 1.5ml of blood was collected from each animal from the femoral vein with 0.5 mL placed in a tube with EDTA for CBC, and the remaining 1.0 ml placed in a serum separator tube for serum chemistry. Blood tubes were transported to SNPRC veterinary clinical pathology core. There samples at the SNPRC were analyzed using the UniCel DxH 800 Coulter Cellular Analysis System for complete blood counts, and the UniCel DxC 700 AU Chemistry Synchron Clinical System was used for serum chemistry panels. Data were scored as either the average or the difference from the nearest post-SSC blood panel subtracting the nearest pre-SSC blood panel reflecting the change in blood chemistry/CBC following the SSCs procedure. One animal was omitted from blood chemistry/CBC analyses for change from post-SSC to pre-SSC as that individual only had a “pre” SSC physical exam/blood work. For a full background on normative ranges of blood chemistry and CBCs in old-aged marmosets, see [37].

### Data Analyses

#### General Analytical Strategy

All data were analyzed using jamovi 2.6 [40] and visualized using Graphpad Prism 10 software or jamovi 2.6. Data were analyzed to assess for the presence of statistically significant mean differences across age, sex, and reactor groups using mixed ANOVAs or t-tests depending on the number of factors. Linear regression analyses were also performed for analyses between different data sets (such as the association between cortisol data and behavioral data). Post-hoc analyses across grouped means in ANOVA were performed and significance was determined using *pTukey* < 0.05. We also performed cluster analyses and cluster plots using the “snowCluster multivariate analysis 7.4.8” add-on in jamovi for performing principal component analysis (PCA) to construct PCA biplots for individual IDs by age, sex, and reactor groups; as well as analyses of hierarchical clustering (creating physiology/behavior dendrograms and behavior/HPA “expression” heatmaps). Data are standardized and clustered by Euclidean distance using the ward.D2 method. Additional data information including descriptive statistics, correlation heat maps of dependent variables, and access to data files are also available in the Supplemental Materials for **SI File 1**. Data were grouped as fixed factors for sex and age group, and for other analyses we included a measure derived from the HPA reactivity data [See Cortisol Data below] to categorically factor reactors and non-reactors, which were differentiated by sample median split by “HPA reactivity” (mean reactor = 76.07 μg Cortisol/mg Creatinine; mean non-reactor = 27.62 μg Cortisol/mg Creatinine) and “HPA reactivity minus baseline” (mean reactor = 40.01 μg Cortisol/mg Creatinine; mean non-reactor = 14.10 μg Cortisol/mg Creatinine) with same individuals split in either case. Typical circadian increase in stress-free marmosets is up to around a maximum peak of 30μg Cortisol/mg Creatinine or 15μg Cortisol/mg Creatinine peak minus baseline [24]. Descriptive t-tests for every measured variable by age, sex, and HPA reactor type is shown in **SI Table 1-3**.

#### Cortisol Data

Urine samples were collected from 6 time points per each SSC (*baseline, hours 1-4, and day-after baseline*). Each of the 14 marmosets underwent 10 SSC sessions resulting in up to 60 urine samples per marmoset. Analyses of the SSC data were collapsed across individual replicates in three ways. **1)** the first two hours of the SSC were collapsed into one time point (by averaging SSC hour 1 and hour 2 cortisol), and the last two hours of the SSC were collapsed together into another time point (by averaging SSC hour 3 and hour 4 cortisol), which resulted in the <u>SSC having four time points</u> (*baseline, first half, second half, day-after baseline*). This was performed to minimize missing data as not every marmoset provided a urine sample every hour of every SSC. Specifically, out of a total number of 840 possible urine samples we aimed to have collected (i.e., *14 marmosets * 10 SSCs * 6 time points*), we had a ∼87% success rate (13% missing) for urine collection. Collapsing to our four time points resulted in having ∼98% of samples (2% missing) across the 14 marmosets’ 10 SSCs. **2)** The SSC data were analyzed with all 10 SSCs collapsed and averaged across time points. In other words, each marmoset had 10 replicates averaged to form an individual SSC response, and these data were analyzed for group differences by age, sex, and reactor status on each of the dependent physiological and behavioral measures on their averaged SSC data (behavioral measures from each of the 10 SSCs were collapsed in the same method). **3)** Lastly, in order to test the hypothesis that individual marmosets habituate to repeated SSC’s, we compared the physiological and behavioral parameters of the first 3 SSCs to the last 3 SSC sessions. Whether analyses were on all 10 SSCs, or first/last 3 SSCs is explicitly indicated in the results.

In addition to the analyses of the raw urinary cortisol data concentrations (µg/mg Creatinine) across the SSCs, we derived multiple variables that reflect different components of HPA activity. <u>HPA Reactivity:</u> Reactivity represents the idea of how much (both in magnitude and duration) an individual marmoset physiologically responds to the SSC stressor event, with higher reactivity suggesting a marmoset was more responsive or reactive to the SSC stressor. HPA reactivity was calculated in multiple ways to reflect different physiological constructs. For each individual SSC, we calculated the maximum cortisol concentration as “REACMAX”; the maximum cortisol concentration – the baseline cortisol concentration as “REACMAX_BL”; the percentage change of the maximum cortisol concentration relative to the baseline concentration as “REAC%”; The area under the curve (AUC) of the individual SSCs where the floor was zero as “AUC”; The area under the curve of the individual SSCs where the floor was the baseline concentration as “AUC_BL” similarly to methods in marmoset stress reactivity published previously [41]. AUC was calculated using the trapezoidal rule (summation of trapezoidal areas across each series of data) in excel. Each of these derived HPA reactivity measures were highly correlated, but each reflects subtle differences in HPA physiology both accounting for basal concentrations of cortisol and magnitude and duration of HPA peak responses during SSCs that may vary by sex, age, or behavioral response. <u>HPA Recovery</u>: Recovery represents the idea of how close the individual marmoset returns to their normative baseline the day following the SSC stressor, with higher recovery values suggesting a marmoset’s day-after baseline was higher than the baseline the day-of the SSC stressor. Higher recovery values may suggest that the marmoset’s response to the SSC had a longer residual effect or remained reactive longer. HPA recovery was calculated by taking the day-after baseline cortisol concentration and subtracting the baseline cortisol concentration the day of the SSC. This value was presented both as the cortisol concentration difference “RECOVERY” and the % difference “RECOVERY%”.

## RESULTS

### SSCs produce an increase in urinary cortisol

The overall change in cortisol over time during the SSCs was significant *F*(3,39)=16.3, *p*<.001, demonstrating that cortisol increased within the SSC period. Specifically, urinary cortisol was higher from the first half period (hours 1-2) of the SSC (*t*(12)=-4.4, *p*=.004) and the second half period (hours 3-4) of the SSC (*t*(12)=-5.03, *p*<.001) compared to day-of SSC baseline. The day-after baseline urinary cortisol concentration was also higher than the day-of SSC baseline (*t*(12)=-3.6, *p*=.011), suggesting the SSC resulted in a residually higher basal cortisol concentration a day after the SSC. The second half urinary cortisol was higher than the first half urinary cortisol during SSCs (*t*(12)=-3.6, *p*=.011). The day-after baseline cortisol was lower than the second half urinary cortisol during SSCs (*t*(12)=-3.9, *p*=.009), but there was no difference between day-after baseline cortisol concentrations and the first-half period of the SSC cortisol concentrations (*t*(12)=1.03, *p*=.734). The averaged pattern of urinary cortisol response during SSCs is shown in **Figure 2**

**Figure 2.**
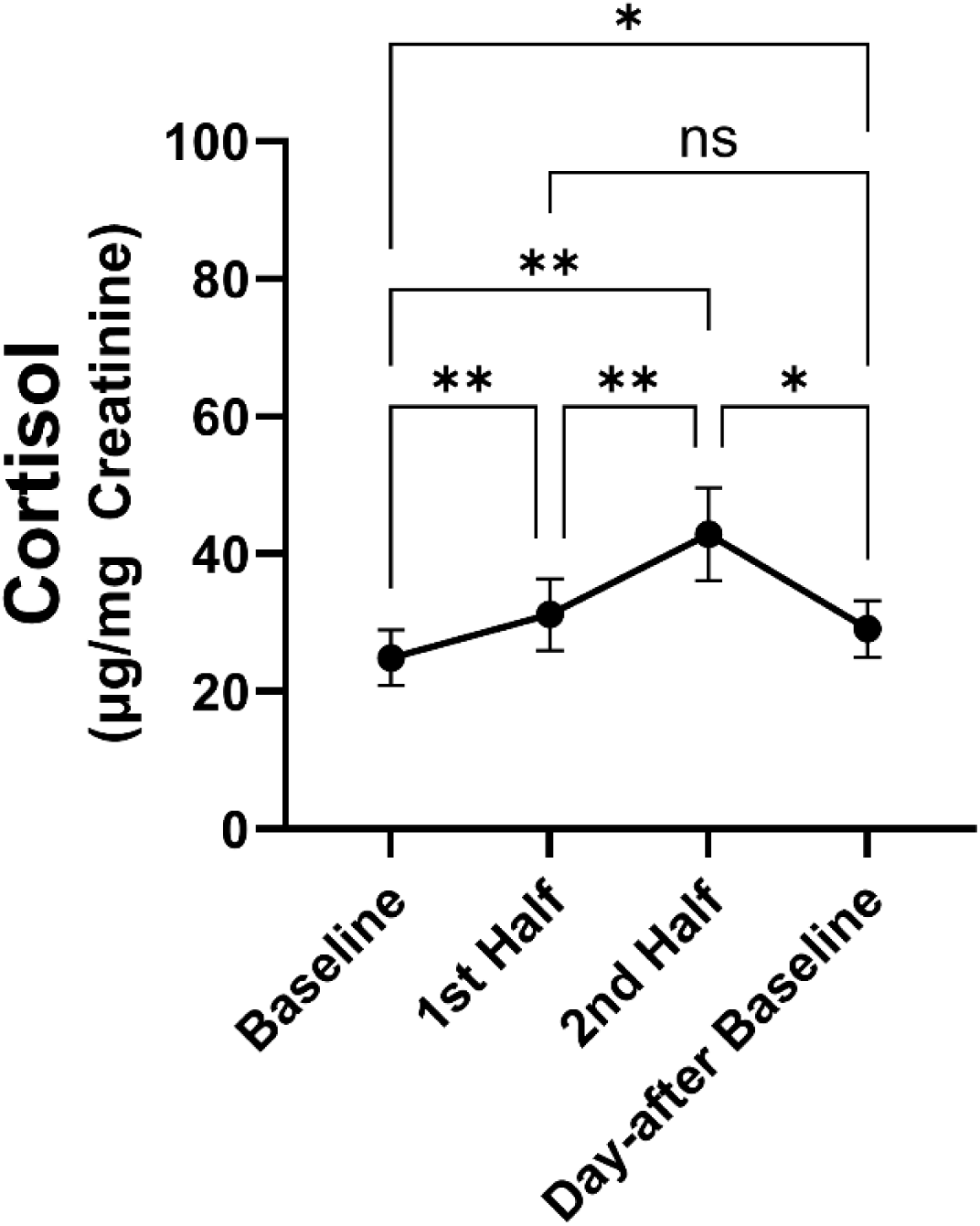
Average urinary cortisol ± SEM across all SSCs for all marmosets. *= *p*<.05; **= *p*<.01; ns = not significant.

### Individual differences in HPA responses to SSCs in aging marmosets: reactors and non-reactors

When examining individual averaged SSC responses, there is a strong appearance of discrete physiological phenotypes in response to SSCs, where some marmosets show high reactivity and others show little to no reactivity (**Figure 3**). Of the 14 marmosets, half of them showed what is a clear increase in urinary cortisol (every HPA reactivity >40μg Cortisol/mg Creatinine) during the SSC, which we have referred to as “reactors”. The other half of the marmosets showed little or no increase in urinary cortisol during the SSC, which we have referred to as “non-reactors” (every HPA reactivity <35μg Cortisol/mg Creatinine). Reactors had higher day-of and day-after baseline values (*t(12)*>4.34, *p*<.001), were higher in each of the HPA reactivity measures (*t(12)*>3.73, *p*<.003), and showed higher recovery% (*t(12)*=2.2, *p*=.048). However, there was no difference between reactors and non-reactors for recovery or reactivity%. There was also no contingency for grouping reactors vs. non-reactors by sex (*χ^2^*=1.17, *p*=.28) or age group (*χ^2^*=2.57, *p*=.10), though there may be a trend for higher number of non-reactors in the very-geri age group (*5 out of 7 very-geris*). A heatmap visualization of the differences in HPA and reunion behavioral parameters between reactors and non-reactors is shown in **Figure 4**. For urinary cortisol SSC plots separated by each individual see **SI Figure 1**.

**Figure 3.**
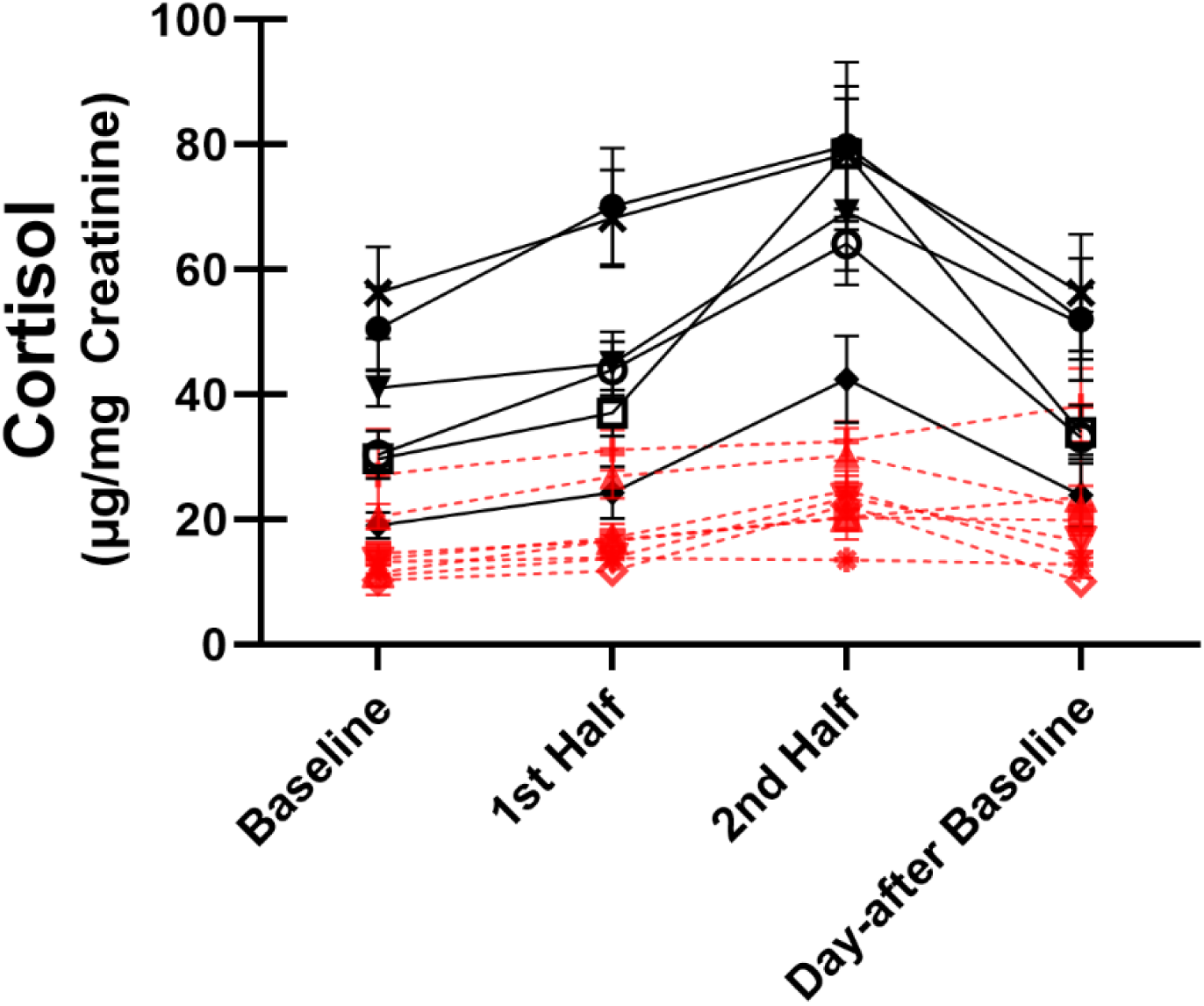
Average urinary cortisol ± SEM across all SSCs. Marmosets are grouped by reactors (black, solid line) and non-reactors (red, dashed line) based on difference in urinary cortisol change (HPA reactivity) during SSC. Each individual line corresponds to the averaged SSC response for an individual marmoset.

**Figure 4.**
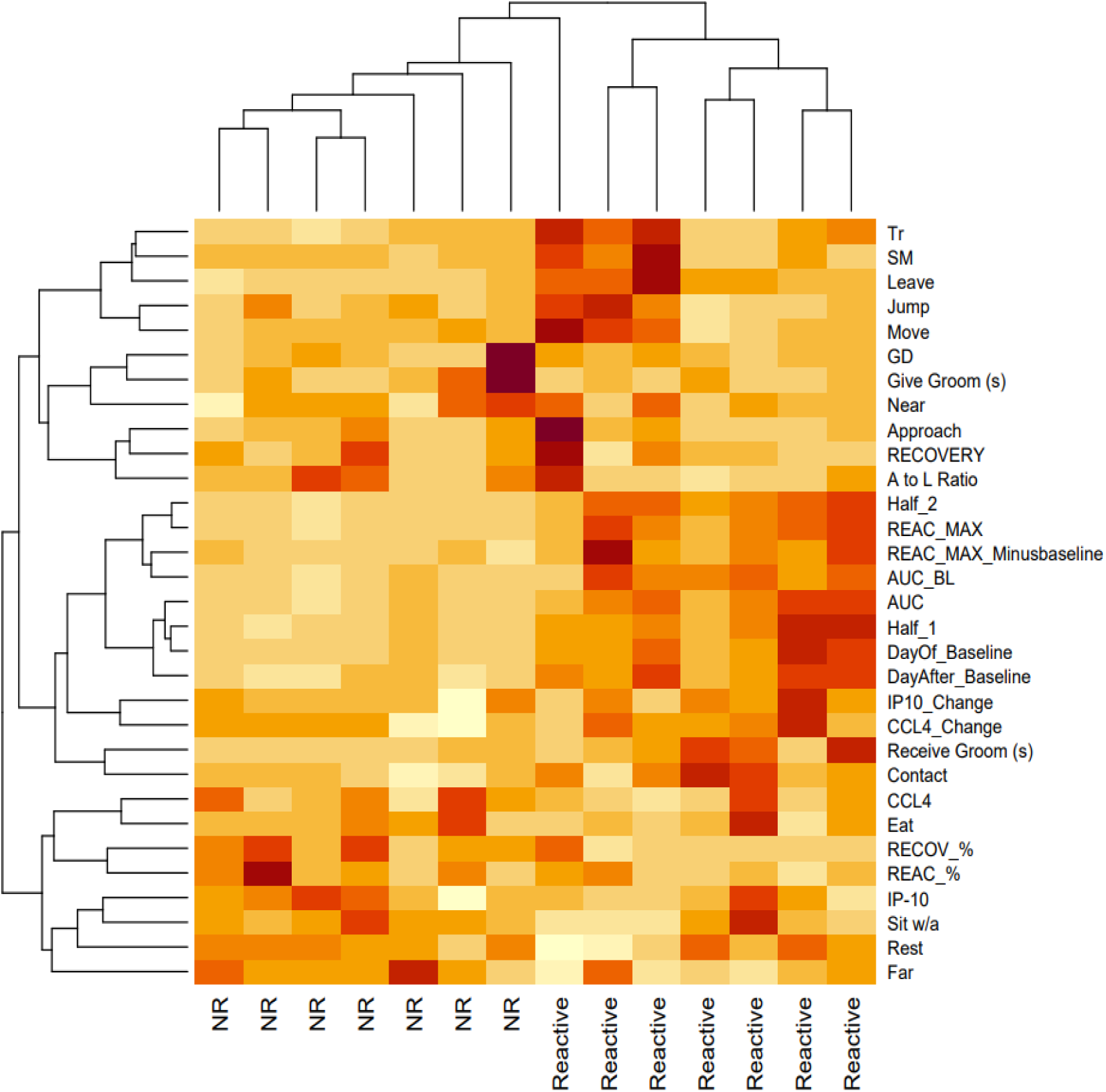
HPA and reunion behavior heatmap clustered by reactor (Reactive) and non-reactor (NR) phenotypes. Each column represents an individual marmoset. Higher color intensity reflects higher value outcome. Tr = move transitions; SM = scent marking; GD = genital display. See methods/Table 1 for full name descriptions.

### Differences in HPA responses by age group and sex

The overall change in urinary cortisol during the SSCs did not differ by age group *F*(3,30)=.875, *p*=.465. Both peri-geri and very-geri marmosets showed the same change in urinary cortisol across SSCs, and did not differ in any of the HPA reactivity or HPA recovery parameters (**Figure 5**). The overall change in urinary cortisol during SSCs did, however, differ by sex *F*(3,30)=4.39, *p*=.011. Females showed higher urinary cortisol during SSCs (**Figure 6)**. While females showed greater change in urinary cortisol during the SSCs compared to males, only HPA recovery% was significantly different (*t(12)*=2.25, *p*=.044), suggesting females were also better at recovering from SSCs. Interestingly, there was a sex by age interaction in the change in urinary cortisol during SSCs *F*(3,30)=3.24, *p*=.036 (**Figure 7**). In general peri-geri females show the highest urinary cortisol concentrations and reactivity, while very-geri females show no difference compared to males. The very-geri females show HPA reactivity responses that appear to converge toward reactivity patterns similar to “non-reactors”. Moreover, females in general show better HPA recovery%, with peri-geri females showing the strongest return to baseline with lower recovery% cortisol compared to peri-geri males (*t*(10)=3.18, *p*=0.04) (**Figure 7**).

**Figure 5.**
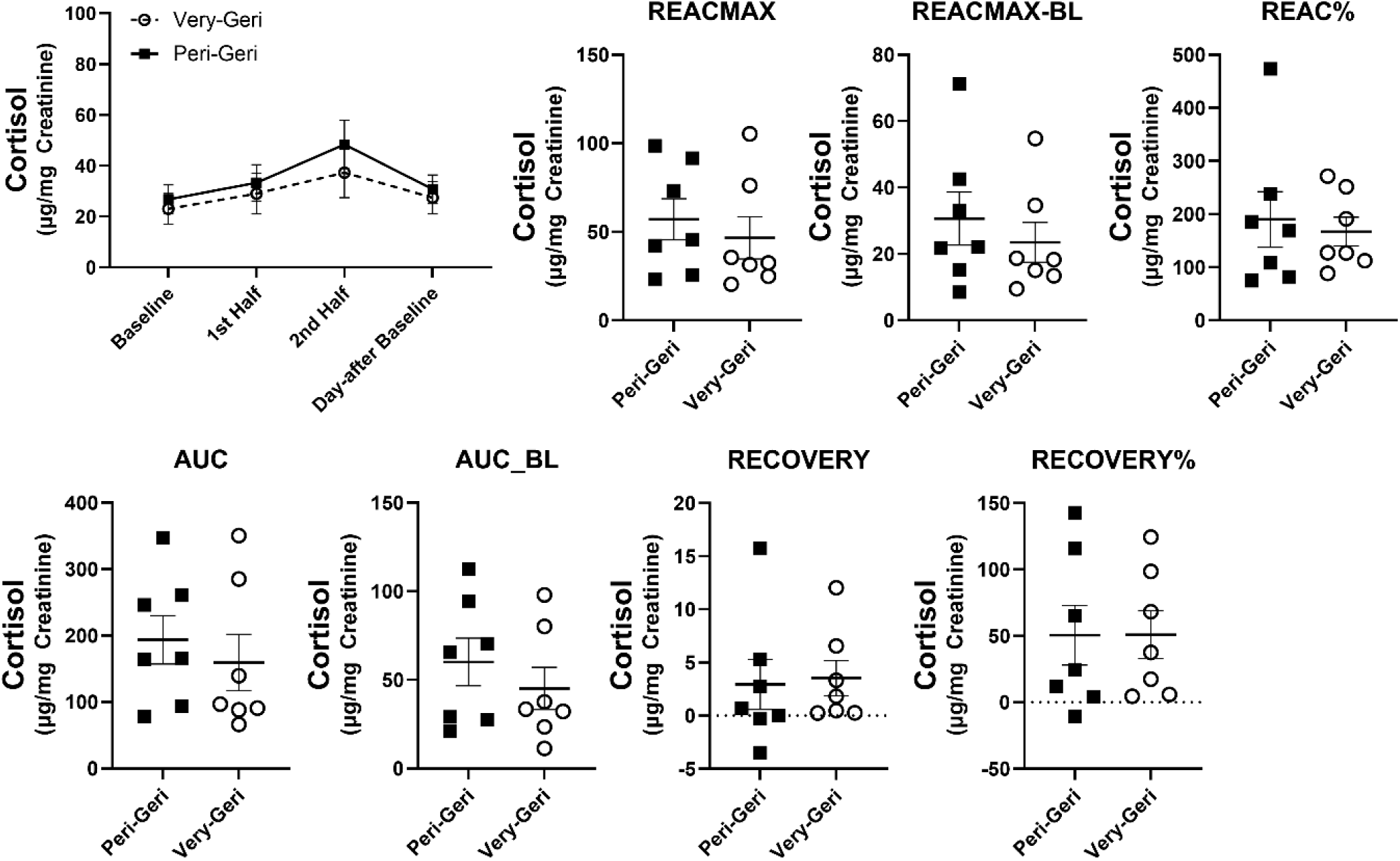
Average urinary cortisol ± SEM across all SSCs by age group (peri-geri and very-geri), and mean differences in urinary cortisol ± SEM in each of the HPA reactivity and HPA recovery parameters by age group. Individual data points reflect individual marmoset averaged responses.

**Figure 6.**
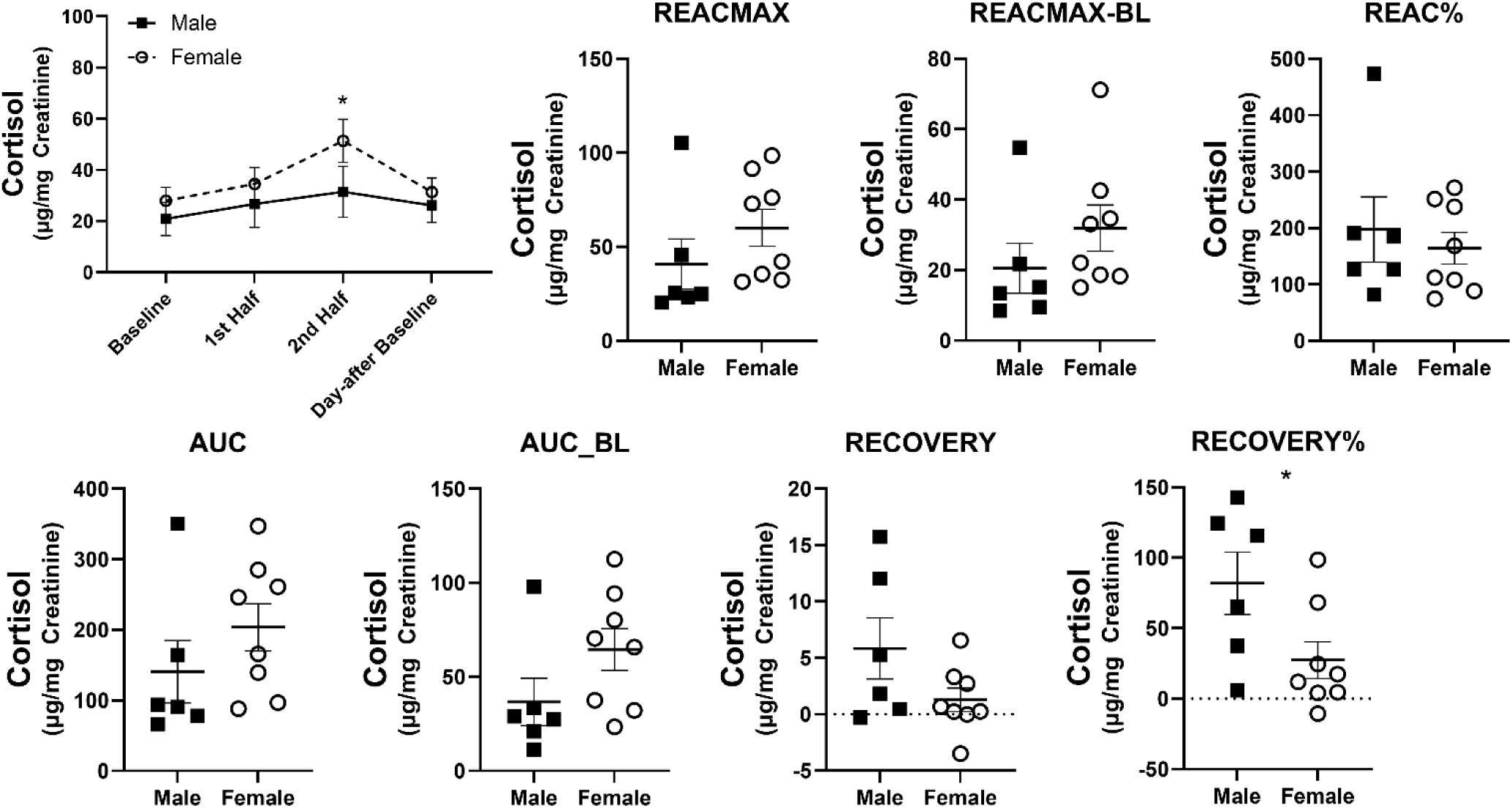
Average urinary cortisol ± SEM across all SSCs by sex. and mean differences in urinary cortisol ± SEM in each of the HPA reactivity and HPA recovery parameters by sex. Individual data points reflect individual marmoset averaged responses. * indicates *p*<.05.

**Figure 7.**
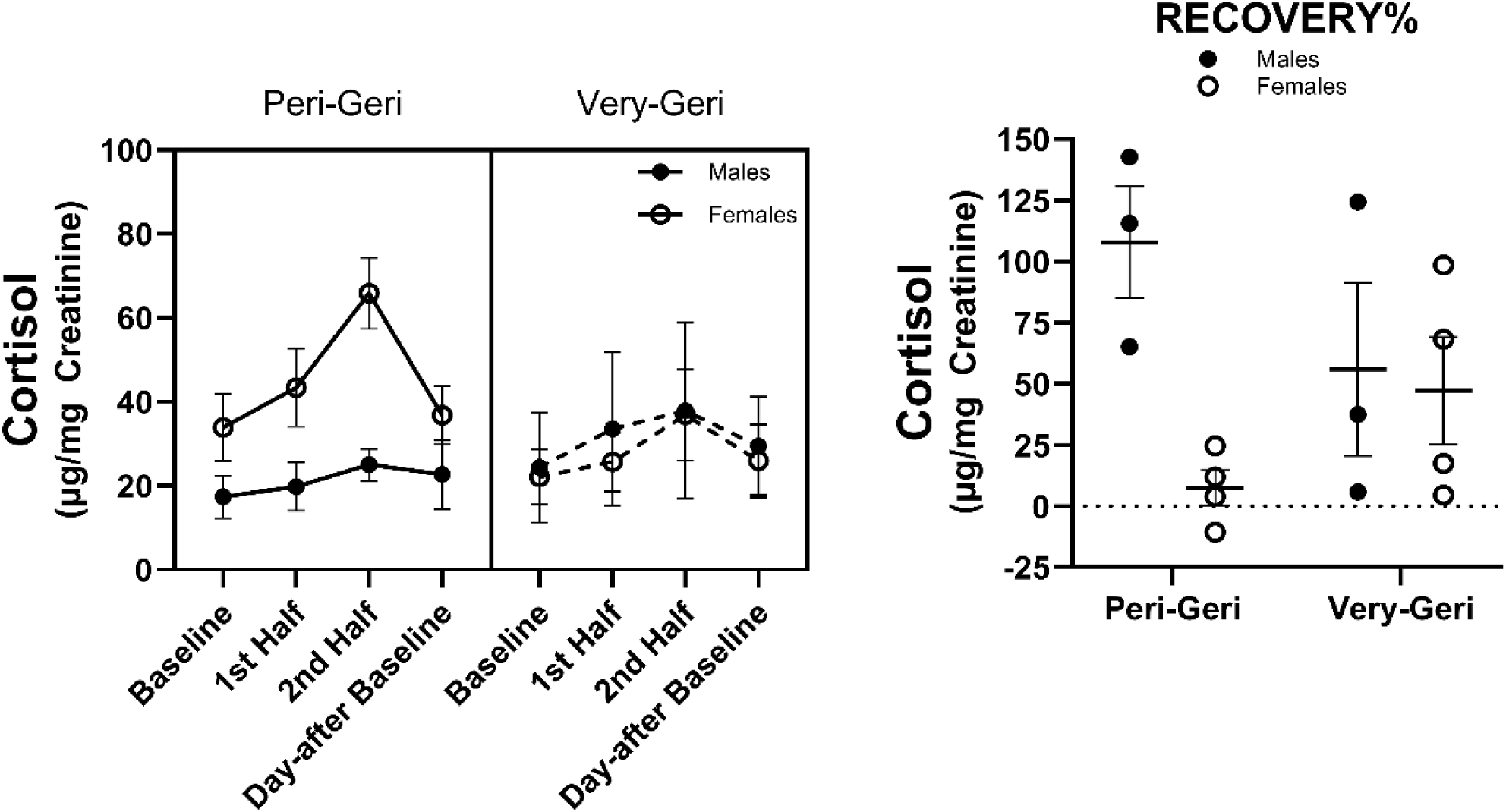
Average urinary cortisol ± SEM across all SSCs by both sex and age group, and mean difference ± SEM in recovery% by both sex and age group. Individual data points reflect individual marmoset averaged responses.

### Habituation of HPA responses across SSCs

We were interested in whether marmosets showed differences in habituation to the repeated SSCs which would be demonstrated by finding that the later SSC sessions have significant differences in urinary cortisol and behavioral responses compared to earlier SSC sessions. We analyzed whether there were changes in outcomes between the first three SSC sessions compared to the last three SSC sessions. Overall, we found that the change in cortisol during SSCs did not vary by session *F*(3,30)=.487, *p*=.694, nor did the change in cortisol during the first and last three SSCs interact with age or sex *F*(3,30)<.81, *p*>.39 (**Figure 8**). There was a trend for a session by age by sex interaction *F*(1,10)=4.09, *p*=.071. The only HPA reactivity, recovery, or behavioral parameters that differed or interacted with age and sex between the first and last three SSC sessions are shown in **Figure 8**. There were interactions in move transitions (*F*(1,10)=13.96, *p*=.004), max reactivity (*F*(1,10)=4.72, *p*=.055), approaches (*F*(1,10)=6.75, *p*=.027), and approach to leave ratio (*F*(1,10)=31.11, *p*<.001), each of which varied between first and last three sessions by age or sex. Correlations between HPA and behavioral measures during the first three SSCs and the last three SSCs are shown in **SI Figure 2**.

**Figure 8.**
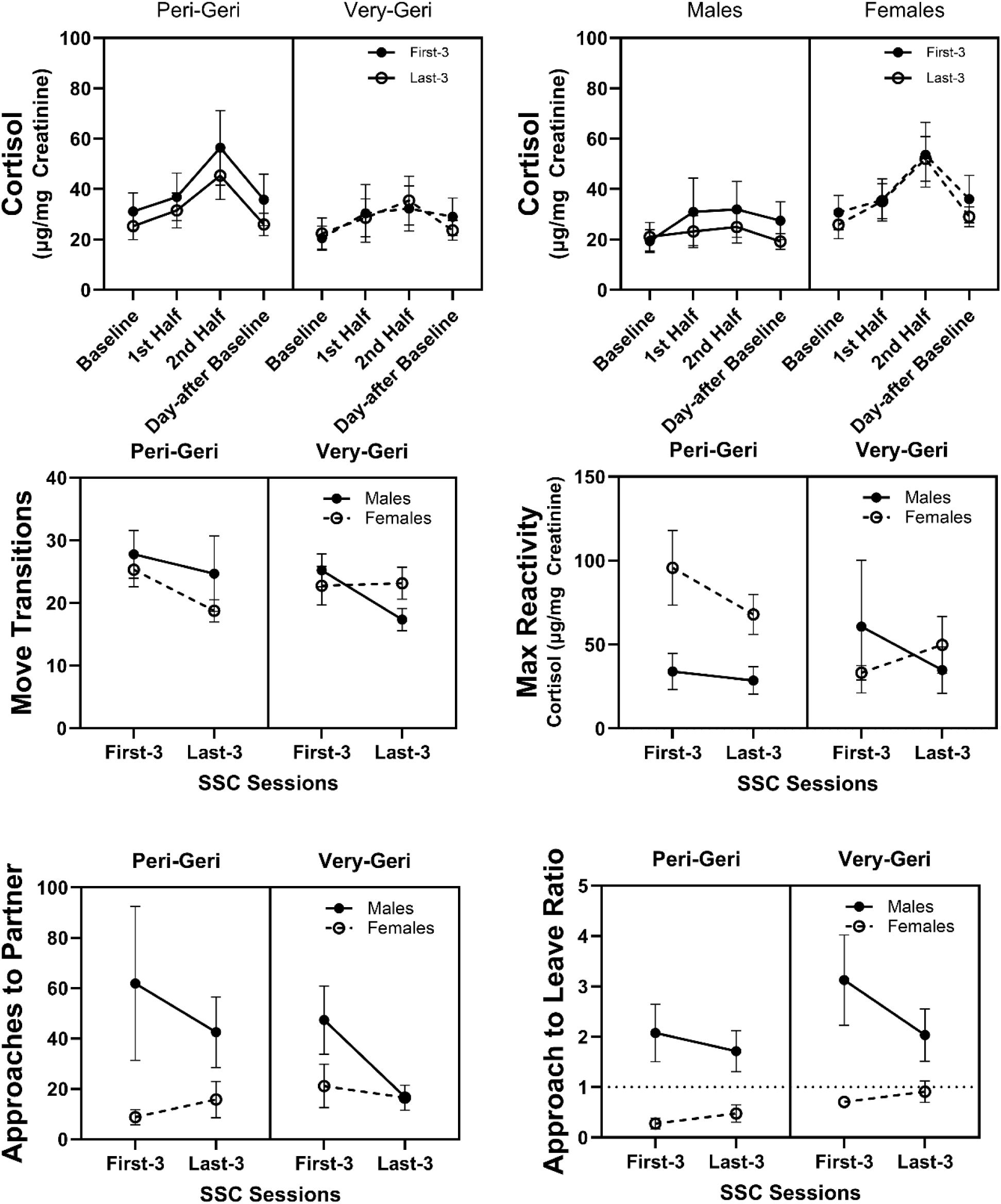
Average urinary cortisol ± SEM across the first three and last three SSCs separated by age group and sex. Averaged move transitions, max reactivity, approaches, and approach to leave ratio each showed significant differences and interaction effects between first three and last three SSCs and sex.

### Correlates of isolation behavior and HPA activity

Overall, behaviors during isolation were not correlated with any HPA activity. Specifically locomotive behaviors such as moving, resting, jumping, sitting, and standing were not associated with measures of either HPA reactivity or recovery (all *r*(12)<.30, *p*>.29). Vocalizations suggesting stress or long-range contact to partners (alarm/phee/shrill calls) also did not correlate with reactivity or recovery (all *r*(12)<.32, *p*>.27). Other behaviors such as head scanning, scent marking, self-grooming, and time spent in their enclosure bucket did not correlate with reactivity or recovery either (all *r*(12)<.21, *p*>.46). Thus, taken as a whole, behavior during the isolation period did not predict HPA activity. The frequency or duration of all isolation behaviors did not differ between peri-geri and very-geri age groups, with only the frequency of head scanning showing a trend to decrease in very-geri individuals *t*(12)=1.98, *p*=.07. However, there are notable overall sex differences in the frequency and duration of isolation behaviors, where rates of jumping (*t*(12)=2.24, *p*=.045), move transitions (*t*(12)=2.73, *p*=.02), and scent marking were higher in females compared to males (*t*(12)=2.02, *p*=.06); while rates of sitting (*t*(12)=2.88, *p*=.01) and eating (*t*(12)=2.00, *p*=.07) were higher or trending higher in males compared to females. Interestingly, reactors and non-reactors showed no behavioral differences in any of the isolation behaviors, corroborating the idea that behavior at the onset of the SSC does not predict HPA activity (all *t*(12)<1.18, *p*>.26).

### Correlates of HPA activity and reunion behavior

We found that HPA reactivity was correlated with a few reunion behaviors. Specifically, both REACMAX and AUC were positively correlated with the number of move transitions in the cage *r*(12)=.564, *p*=.036. Locomotion (move transitions) was also correlated with many other parameters including day-of SSC baseline (*r*(12)=.579, *p*=.03), day-after SSC baseline (*r*(12)=.701, *p*=.005), jumping (*r*(12)=.723, *p*=.003), approaches (*r*(12)=.667, *p*=.009), and leaves (*r*(12)=.828, *p*<.001), and negatively correlated with sitting *(r*(12)=-.716, *p*=.004). AUC was correlated with the duration of received grooming *r*(12)=.537, *p*=.048. Lastly, AUC (*r*(12)=.504, *p*=.066) and REACMAX (*r*(12)=.564, *p*=.07) both showed a positive trend for correlation with the frequency of leaving their partner during reunion. However, HPA reactivity was not correlated with other important reunion behaviors including time spent in contact with or nearness to their partner, approaching partners, duration of sitting, or frequency of jumping. HPA recovery showed a different profile of correlations with reunion behavior. Specifically, recovery was positively associated with both the frequency of approaching their partner (*r*(12)=.815, *p*<.001), and higher “approach to leave” ratios (*r*(12)=.714, *p*=.004), both suggesting that marmosets who more frequently approached their partner had higher recovery values. Lastly recovery% was borderline negatively associated with “received grooming” (*r*(12)=-.528, *p*=.052), suggesting animals who received more grooming had an overall closer percent return to normal baseline the day after their SSCs. A summary of these correlations is shown in **Figure 9**. Overall, the rate or duration of reunion behaviors did not differ by age group (all *t*(12)<1.61, *p*>.13), but some behaviors differed by sex including the frequency to approach their partner (*t*(12)=2.26, *p*=.04), the approach to leave ratio (*t*(12)=5.03, *p*<.001), and a trend in genital displays (*t*(12)=1.88, *p*=.085), each of which was higher in males compared to females. An overall summary of these data is shown in **SI Figure 3**.

**Figure 9.**
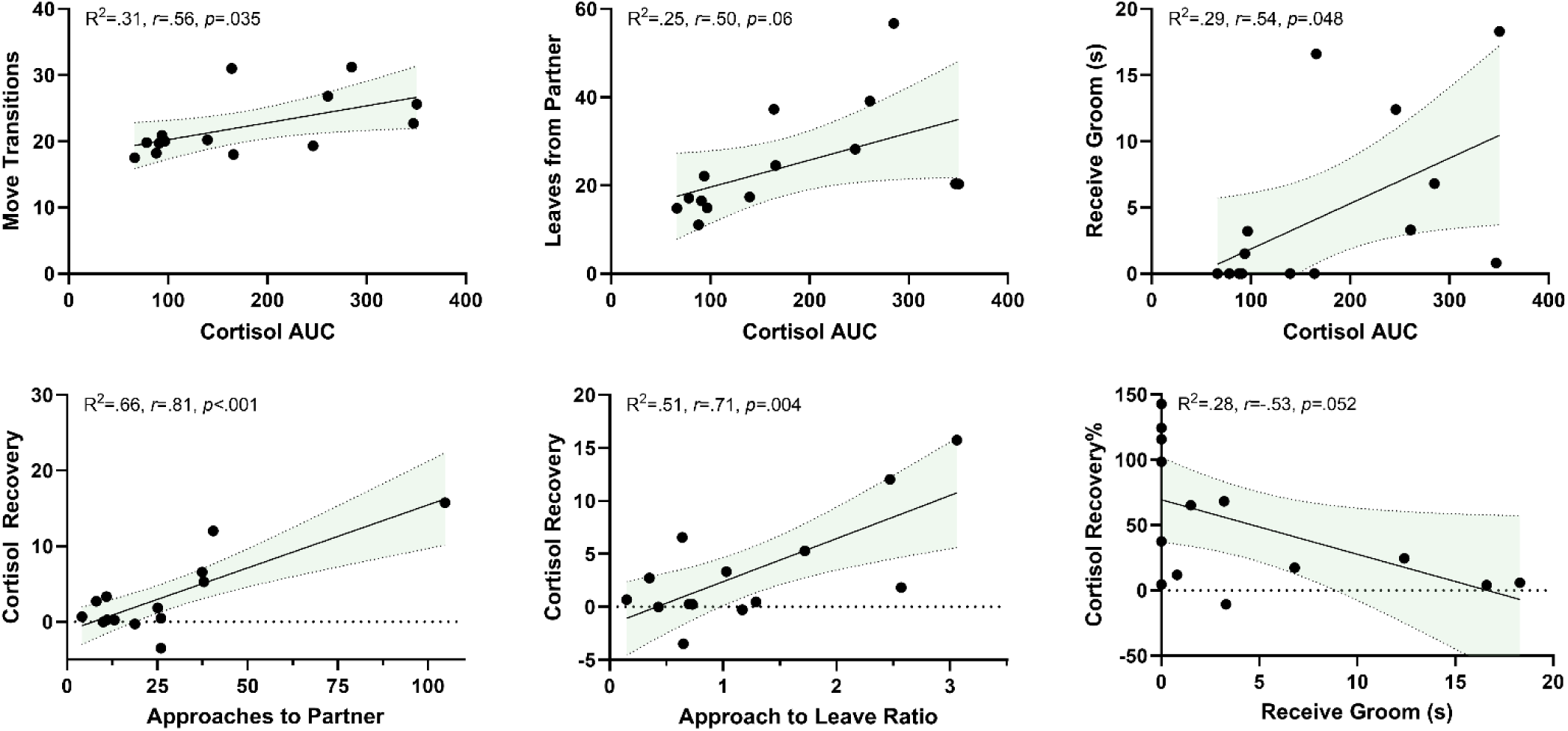
Representative significant correlations between cortisol AUC and cortisol recovery with reunion behavior parameters. Parameters presented on the X-axis are outcomes that precede the outcomes presented on the Y-axis (i.e., *cortisol reactivity occurred before social reunion and social reunion occurred before cortisol recovery*). Shaded area represents 95% confidence intervals.

### Correlates of HPA activity with body weight change, blood chemistry, inflammatory markers, and CBCs

Specifically examining which parameters of blood chemistry and CBCs were correlated with HPA reactivity or HPA recovery, we found two parameters that had strong relationships with HPA reactivity and recovery. We found that the change in blood glucose following SSCs was positively correlated with day-of SSC baseline (*r*(11)=.565, *p*=.044), day-after SSC baseline (*r*(11)=.585, *p*=.036), REACMAX (*r*(11)=.603, *p*=.029), AUC (*r*(11)=.585, *p*=.036), and negatively associated with recovery% (*r*(11)=-.698, *p*=.008). Moreover, neutrophil % cell counts were negatively associated with REACMAX (*r*(11)=-.615, *p*=.025), AUC (*r*(12)=-.509, *p*=.075), and day-of SSC baseline (*r*(11)=-.499, *p*=.082). The change in body weight was not correlated with any HPA parameters, but the change in weight over the 6 month period from the start of the SSCs was positively correlated with the change in blood glucose (*r*(11)=.553, *p*=.050), and the maximum weight change over the year was positively correlated with the change in cytokine/chemokine markers CCL4 (*r*(12)=.597, *p*=.024) and CXCL10 (*r*(12)=.635, *p*=.015). A summary of these correlations is shown in **Figure 10**. A correlation matrix of all parameters is shown in **SI Figure 4**.

**Figure 10.**
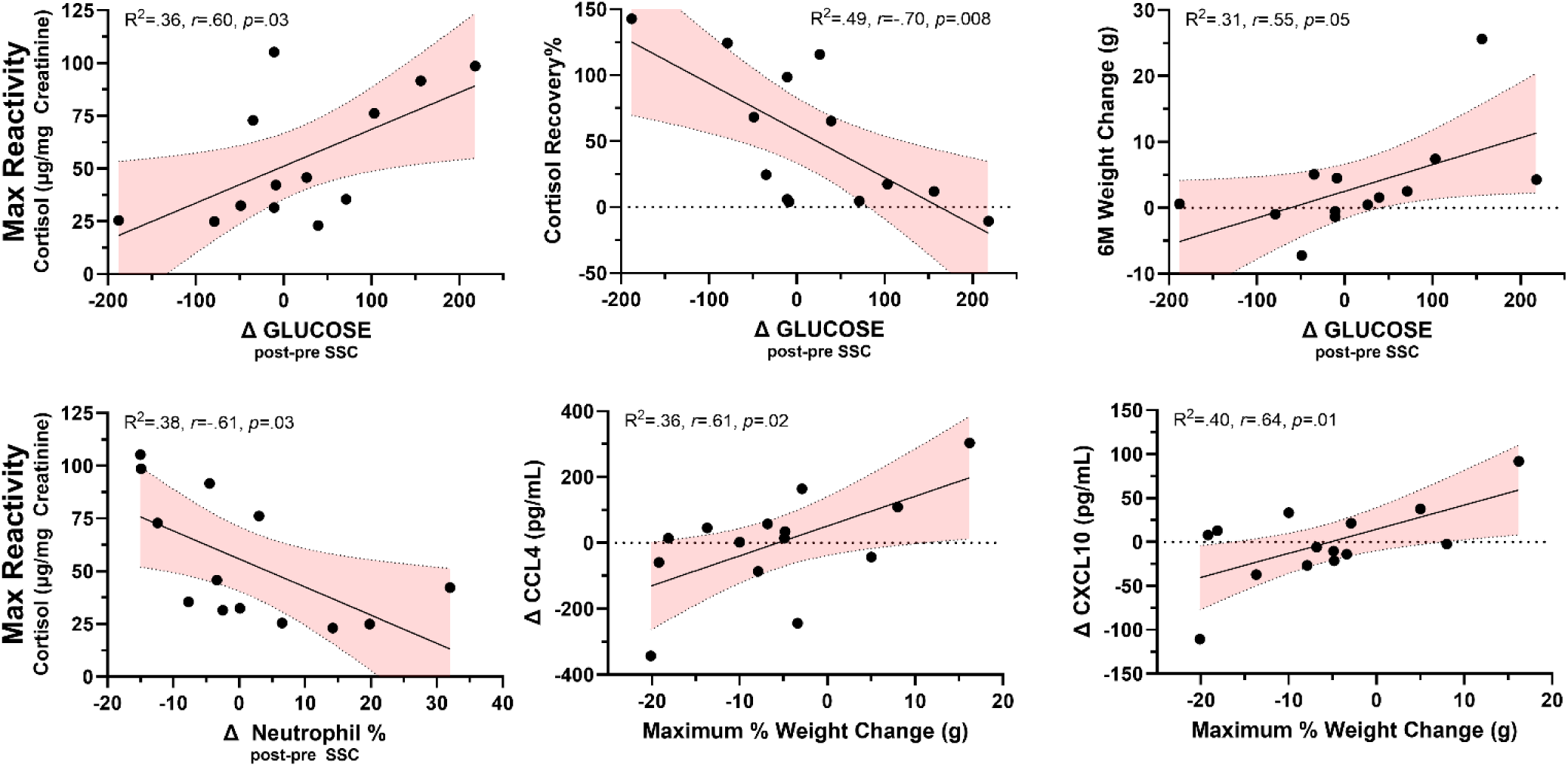
Representative significant correlations between HPA reactivity and recovery and health parameters. 6M Weight change is the weight change (g) from the beginning of the SSC sessions and 6 months after the start of SSCs. The maximum % weight change reflects the largest observed weight change by % measured over the course of the recent year during SSCs. Change in glucose and change in neutrophil cell count % reflect post SSC – pre SSC values. Conversely, the change in cytokine/chemokine CCL4 and CXCL10 reflect the change in plasma concentration between a sample collected at the start of the SSC and the sample collected at the end of the SSC (i.e*., positive values reflect an increase in cytokine concentration over the SSC period*). Shaded area represents 95% confidence intervals.

### Differences between reactors and non-reactors in health parameters and reunion behavior

Given the clear physiological difference in some marmosets not showing a strong physiological response to the SSCs, we were interested in investigating whether there are negative health outcomes associated with reactor or non-reactor SSC response phenotypes. We evaluated differences in both health parameters and behavioral responses to see which outcomes differed between reactor and non-reactor groups. We found that non-reactors had reduced weight gain 6 months post-SSCs *t*(12)=-2.3, *p*=.04, higher total calcium *t*(12)=2.26, *p*=.043, higher change in calcium *t*(11)=2.25, *p*=.046, higher change in phosphorus *t*(11)=2.19, *p*=.052, and higher change in white blood cell counts *t*(11)=2.20, *p*=.050, shown in **Figure 11**. With regard to reunion behavior outcomes, non-reactors showed fewer move transitions *t*(12)=-2.69, *p*=.02, reduced time spent in contact with their partner *t*(12)=-2.57, *p*=.024, reduced number of times leaving partner *t*(12)=-3.15, *p*=.008, and received less grooming from their partner *t*(12)=-2.68, *p*=.02, shown in **Figure 12**. Overall reactors and non-reactors appear to show distinct phenotypes based on components of HPA activity and behavior **SI Figure 5**.

**Figure 11.**
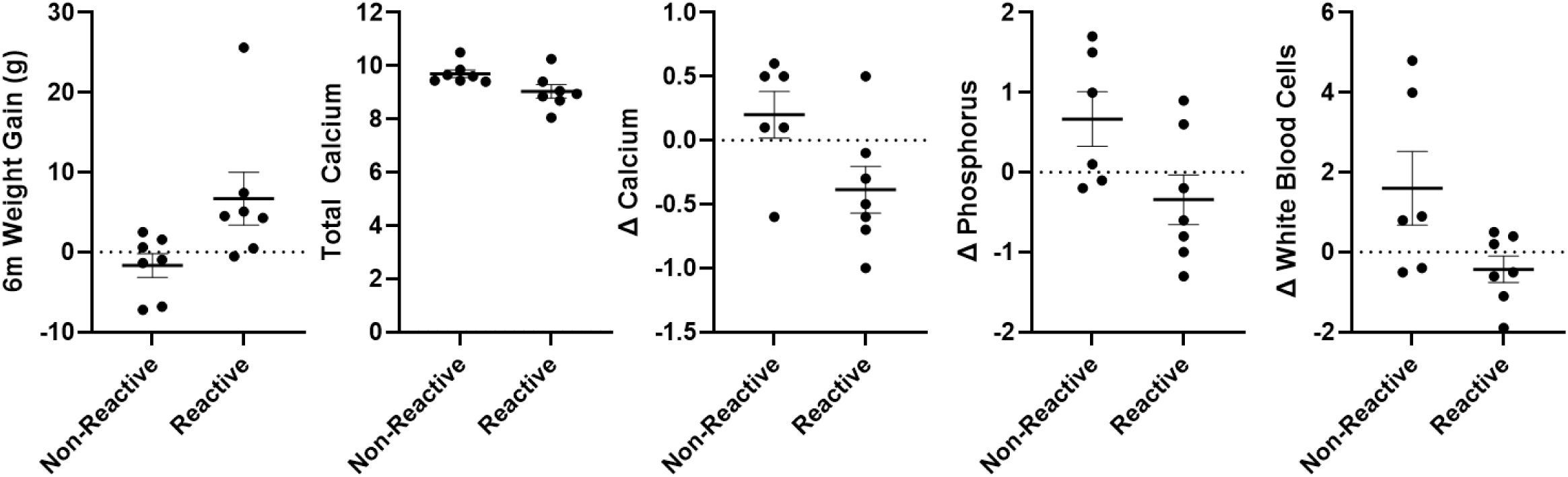
Each panel shows a significant difference (mean ± SEM) in health parameter outcomes between reactor and non-reactor groups. 6M Weight change is the weight change (g) from the beginning of the SSC sessions and 6 months after the start of SSCs. Individual data points reflect individual marmoset averaged responses. Change in blood chemistry and CBC markers reflect post SSC – pre SSC values.

**Figure 12.**
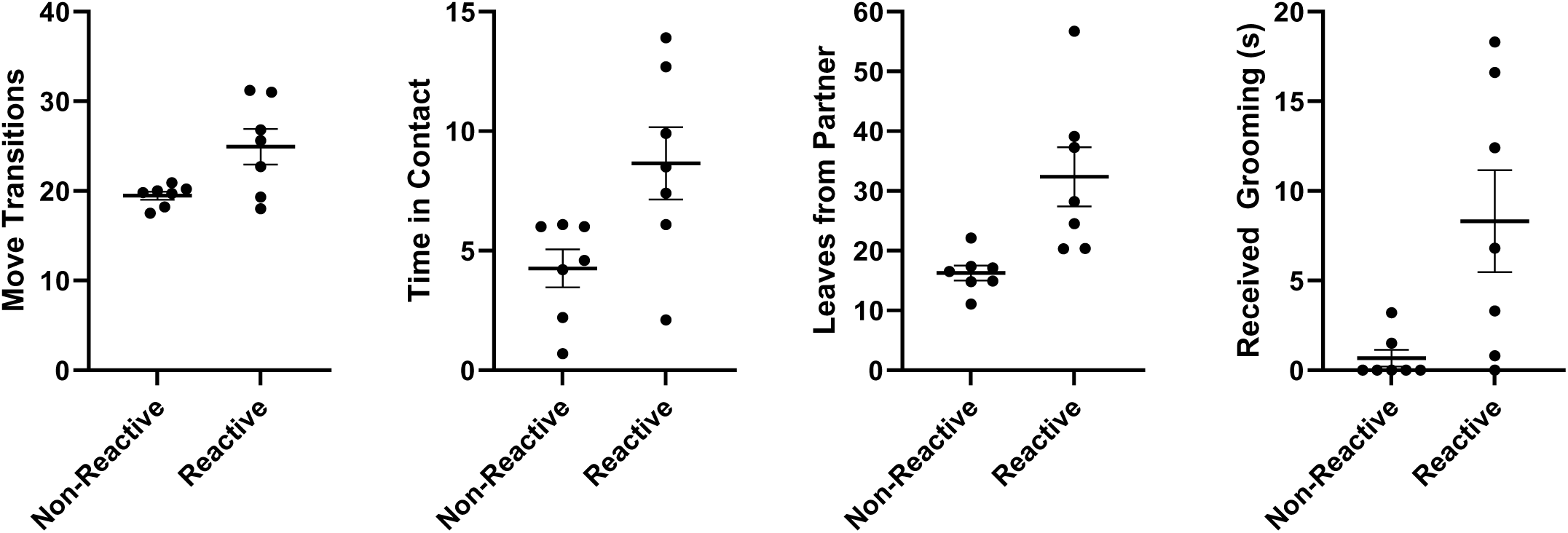
Each panel show a significant difference (mean ± SEM) in reunion behavioral outcomes between reactor and non-reactor groups. Individual data points reflect individual marmoset averaged responses. Behaviors are scored as average per 20 min observation (either in frequency or duration in seconds).

## DISCUSSION

Studying social separation stress in aging marmosets translates to human health by providing insights into the biobehavioral impacts of social isolation in aging, which are relevant to understanding human health and stress-related disorders, depression, and anxiety. Studies have shown that marmosets who experience chronic social stress show behavioral and neuroendocrine changes similar to those observed in depressed humans, including altered cortisol levels, increased anxiety-like behaviors, and anhedonia [42]. Marmoset models help identify the biological pathways involved in stress responses that are linked to physical and mental health risk in humans. Social separation stress studies in aging marmosets also reveal how social support and reunion behaviors modulate recovery from stress, emphasizing the importance of social relationships in resilience during aging. Marmosets are therefore a valuable translational model for developing and testing treatments for human psychiatric conditions in aging.

Overall, we found that responses to SSCs varied considerably depending on multiple factors relevant to both aging and health. The goal of this project was to assess whether aging marmosets showed differences in HPA activity or reunion behavior; and, if so, to identify what physiological or behavioral factors may underlie or mitigate these outcomes. We found that marmosets showed two distinct HPA profiles with roughly half of the marmosets showing normal, adult-like HPA responses [24, 27–28] to the SSCs (reactors), and the other half of marmosets showing little or no HPA reactivity response to the SSC (non-reactors). Female marmosets, especially peri-geri females, had higher HPA reactivity and better HPA recovery than male marmosets, but this difference diminished in older, very-geri marmosets. The overall magnitude of HPA activity was correlated with reunion behaviors, especially locomotive behavior and grooming. However, marmosets who approached more and were groomed less showed less positive HPA recovery and their cortisol remained higher the day-after their SSC. Importantly, we found that reactors and non-reactors showed different profiles of correlations with health parameters such as weight gain/loss, blood chemistry, and metabolic and inflammatory markers. In particular, we found strong positive correlations between blood glucose and HPA reactivity, and strong negative correlations with neutrophil cell counts and HPA reactivity. Lastly, marmosets with reduced HPA activity (i.e., non-reactors) had less weight gain, elevated levels of calcium, phosphorus, and white blood cell counts and reduced behavioral affiliation (less time in contact with their partner and reduced grooming) compared to those with normal HPA reactivity in response to stress (reactors). Overall, old-aged marmosets who show reduced HPA responses to social stressors may have different vulnerabilities to negative health and behavioral outcomes, and males appear more likely to to present with this HPA non-reactor phenotype earlier in age.

To date, only a small number of studies have explicitly tested social separation stress in older aged marmosets. One such study found that on average, older marmosets showed increased HPA reactivity relative to baseline [43] similarly to what you would find in younger adult marmosets [24, 27–28]. The marmosets in this study are near in age to our peri-geri age category (5 to 7 years old) and, while not included as part of the analyses in that study, the data provided show that some individuals in their sample also presented with little or no change in cortisol reactivity during the stressor, which is similar to our findings in our older marmoset sample here. These HPA activity patterns found among older marmosets were stable across time on average, but it is uncertain from the data whether individuals who showed no HPA reactivity in this study during year one showed the same lack of reactivity during later years [43]. Interestingly, Rothwell et al., found no sex differences in either HPA reactivity, HPA recovery, or in behavioral responses during testing, but it is worth noting that females in this cohort did show greater cognitive decline compared to males [43]. Whether altered HPA activity was correlated with cognitive performance was not tested, but this would be an interesting area of future investigation in marmosets. Other work has shown a negative correlation with hair cortisol levels and age in marmosets, though notably the differences between aged and very-aged marmosets were not significant [44], with most of the effects being attributed to differences between adult and juvenile/infant marmosets. Additionally, female marmosets both in captivity [24, 44–45] and in the wild [46] had higher cortisol concentrations compared to males, and greater responsiveness in a dexamethasone challenge [47] though sex differences in basal concentrations or HPA reactivity are not always present [48–50]. While many other non-human primate species show age-related changes in HPA activity in response to stressors [51–57], these effects are frequently understudied in aging populations, as many of these studies only examined age ranges of adults rather than geriatric populations. It is worth noting, though, that outcomes in very-geri marmosets may also reflect survivor effects, i.e., the healthiest animals live to very-geri ages in marmosets. While cross-sectional approaches like ours offer important insights, it is important to also assess whether early changes in HPA function may predict poor health outcomes longitudinally.

The impact of hyper or hypo-active HPA activity during aging is important for health. High basal levels of glucocorticoids and the loss of circadian rhythm are known risk factors for cognitive decline and other pathophysiology during aging in humans [58], and excessive activation of the HPA system in the case of chronic stress can have many deleterious effects on nearly every bodily system [59–61]. One of the goals of incorporating repeated social stressors in our study design was to evaluate whether there were age-specific differences in the ability of individuals to habituate or show desensitization to the stressor due to familiarity. While overall HPA activity in response to SSCs did not show signs of habituation or exaggeration during the course of repeated exposure, there was evidence that a few parameters from our findings including max reactivity and social reunion behavior (i.e., locomotion and approaches to partner) did vary in sex and age-specific ways over the duration of the repeated SSCs. While the overall picture of sex and age-related differences in human and non-human primates is very mixed and context-specific, it is important to consider the role of other sex steroid hormones in modulating aging health of the HPA axis and its impact on health, immune function, and cognition [62–64]. One specific caveat in our study is that we didn’t measure changes in testosterone or estradiol in marmosets, which both interact to regulate HPA activity, often in different directions [reviewed in 65]. Unlike humans, female marmosets do not go through menopause and estradiol doesn’t dramatically decline into old age [66]; while male marmosets, like humans, do show a degree of decline in testosterone levels into old age [67–68] Thus, sex-steroid modulation of HPA activity may underlie sex differences in HPA reactivity, HPA recovery, and social behavior during stress.

While our evidence suggests that there are clear differences in a variety of important metabolic and health outcomes associated with the HPA reactor and non-reactor phenotypes in aging marmosets, it remains unclear whether these are definitively positive or negative health outcomes. Previous research has demonstrated that the loss of HPA resiliency or the ability to return or recover back to normal baseline values after a stressor varies by age, sex, and a variety of psychiatric and metabolic diseases [69–72]. In our study, females appear to be more reactive than males, but in our younger, peri-geri group, females show stronger HPA resilience, while males do not; on the other hand, in our very-geri group, the max reactivity is lower and recovery or resilience is weaker in females, which is comparable to very-geri males. What is potentially interesting about these effects is that there was a significant relationship between HPA activity and body weight with important metabolic and immune markers including the change in blood glucose and neutrophil counts. Specifically, observed increases in glucose over the SSC period was positively associated with marmosets who would be considered HPA resilient, (i.e., they are more reactive but recover more strongly) and with marmosets who had higher weight gain (or no weight loss). With regard to immune markers, marmosets who had higher numbers of neutrophils and higher CCL4 and CXCL10 levels over the SSCs also had higher HPA reactivity and larger weight changes. The role of increased CCL4 and CXCL10 concentrations have been implicated in the progression of metabolic diseases or dysregulation including diabetes [73], obesity, and insulin resistance [74–75]; however, to the best of our knowledge no studies have examined these relationships between HPA activity, metabolic health, and chemokine levels in older-aged populations of any species. Overall, it is important to identify whether HPA activity during aging has any bias toward anti-inflammatory or pro-inflammatory regulation of these factors, and future work is needed to disentangle the potential relationship between HPA activation, chemokine levels, and changes in metabolic health in marmosets.

We also noted that calcium and phosphorus concentrations differed between reactors and non-reactors with reactors having decreased levels of both. Previous work has shown that both calcium and phosphorus levels decrease with age, but these effects are sex-dependent with females showing higher declines than males [76]. Interestingly, non-reactor individuals in our study showed an increased change in both calcium and phosphorus levels during the SSC period compared to reactors. Given there were no differences in diet and no effect of menopause, as marmosets do not show human-like menopause [66], these levels could reflect differences in either age-related thyroid function, bone health or kidney function, with elevated phosphorus being a particular risk factor for increased mortality [77–78]. However, in this study, any mechanisms linking HPA activity and calcium and phosphorus levels are unclear and warrant further investigation. Even though we found age-related differences in blood chemistry and CBCs, it is important to consider that these values remained within previously observed age-matched ranges [37].

While this study presents novel data on how aging marmosets respond to repeated social separation stress, there are multiple limitations that should be considered. The first limitation is a small sample size. Though the sample size is small, the point that we tested individuals ten times is a strength that should provide confidence that these physiological and behavioral responses are reflective of age-specific profiles. We also chose deliberately to analyze age as a categorical factor due to the distinction between aging as the transitioning into old age and aging as the state of already being old-age; however correlational data are also included with age as a continuous predictor in supplemental analyses. In general, age as continuous predictor was not strongly correlated with most parameters. Another limitation is that our SSC was only four hours in duration, and it is alternatively possible that non-reactor marmosets could be “slow-reactors” meaning that if we had used a longer separation challenge that we may have seen delayed HPA activation. While unlikely, as most marmoset studies have demonstrated rapid HPA activation [24, 27–28, 43], we cannot rule out this possibility. Another possible explanation for HPA activity differences could be related to previous experimental experience associated with age. Marmosets used in this study were not experimentally-naïve and have variable experimental histories. Another limitation is we did not use a young-age comparison for this study, though these previous studies [24, 27–28, 43] serve as adult normative marmoset responses to social separation challenges for qualitative comparison. Additionally, many of the health parameters were not collected during each separation challenge, and only represent pre- vs post-SSC data; thus, we are correlating data across multiple time scales with behavior, urinary cortisol, and blood chemistry, CBCs, and plasma cytokine/chemokines. With respect to plasma cytokines, there may have been some known stress markers that became elevated after separation but returned to basal level when blood samples were obtained. Lastly, we anticipated that behavioral responses during either isolation or reunion would strongly predict HPA reactivity or recovery respectively. While we found a number of positive associations between reunion behaviors and HPA activity, behavior during the initial isolation period did not correlate with HPA activity or differ by age or reactor/non-reactor type. An important caveat with regard to reunion behavior specifically, is that when an individual is undergoing an SSC in another room, their partner is also alone in the homecage, which may impact their behavior as well [45,50]. Overall with regard to the behavior, there are strong normative sex differences in how marmoset’s respond or perceive social separation as indicated by differences in both reunion and isolation behaviors between males and females.

In conclusion, this study demonstrates a clear distinction between individuals with reactive and non-reactive HPA activity in aging marmosets that are strongly related to multiple health parameters. Moreover, this study shows that sex and age interact as important contributors to differences in physiological and behavioral responses to social stressors. Future work following the impact of these distinct relationships between HPA activity and age/sex should address two key areas: 1) Evaluate whether these differences in HPA phenotypes are associated with other health parameters including more direct measures of cognition, inflammation, gut health, and metabolic health; and 2) examine how the interaction between other signaling systems such as oxytocin and vasopressin, corticotropin-releasing hormone (CRH), and sex-steroids yield biases for either anti- or pro-inflammatory pathways and impact on HPA activity and behavioral outcomes in old age.

## ACKNOWLEDGMENTS

We would like to thank the marmoset team at SNPRC, led by Paulina Castillo, for their excellence in husbandry and marmoset care and welfare, and we would also like to thank Kathy Brasky and their team for outstanding veterinary care of marmosets in this project.

## FUNDING SUPPORT

funding support for this project includes NIH-U34AG068482 awarded to C.R./A.B.S., NIH-R01AG065546 awarded to C.R., NIH-R24OD030215 awarded to L.G., and the SNPRC/Texas Biomed NIH-P51OD011133 and UTHSCSA Pepper Center NIH-P30AG044271.

## Notes

### Competing Interest Statement

The authors have declared no competing interest.

