## Supplemental Materials for "Aging Leads to Altered Physiological Reactivity in Response to Repeated Social Separation Stress in a Nonhuman Primate Model"

**TABLE OF CONTENTS**

SI Figure 1. Plots of average cortisol ± SEM across all ten SSCs for each individual marmoset subjects
SI Figure 2. Correlation heatmap for HPA and behavioral measures during the first and last three SSCs
SI Figure 3. Correlation heatmap for HPA and social reunion behavior for all SSCs
SI Figure 4. Correlation heatmap for Blood Chem/CBC parameters.
SI Figure 5. PCA Biplot from HPA and social reunion behavior parameters for reactor and non-reactor
SI Table 1. T-Tests for each measured parameter based on age group
SI Table 2. T-Tests for each measured parameter based on sex
SI Table 3. T-Tests for each measured parameter based on reactor group
SI File 1. Excel tables of "raw" HPA and behavior data and data averaged across all 10 SSCs.

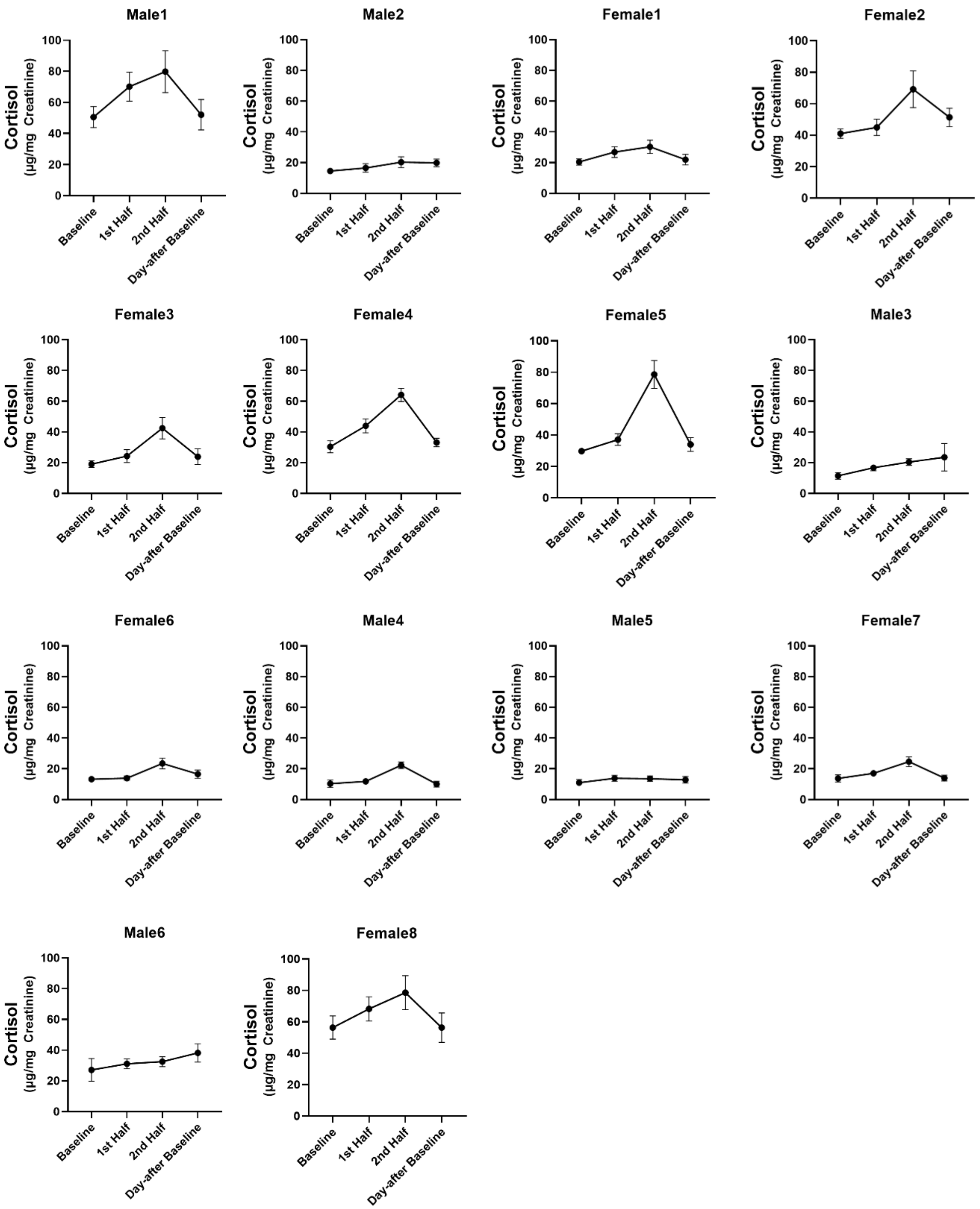


**Figure S1**. Individual plots of cortisol concentration (mean +/- SEM) averaged across all 10 SSC sessions for each individual participant.

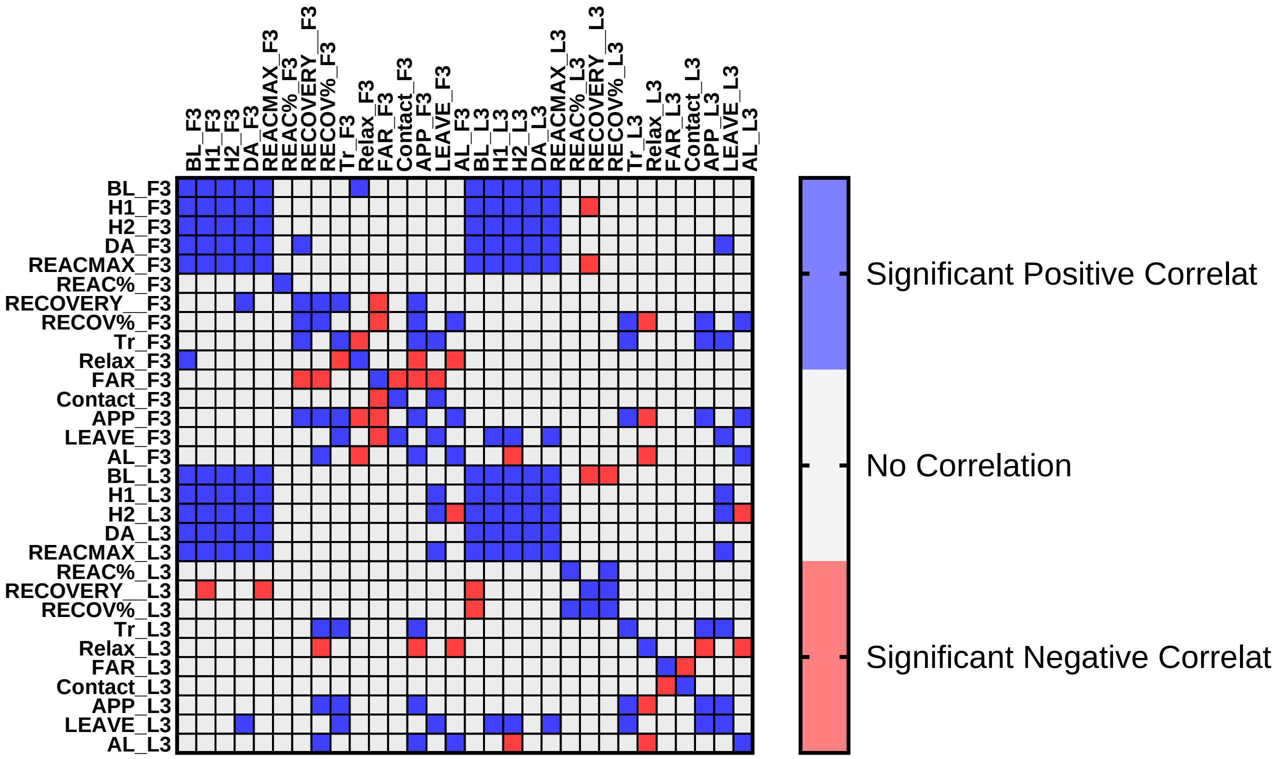


**Figure S2**. Correlation heat map for representative HPA and social reunion parameters during the first three SSCs (F3) and last three SSCs (L3). APP = approaches, AL = approach to leave ratio. BL = baseline cortisol; DA = day-after baseline cortisol.

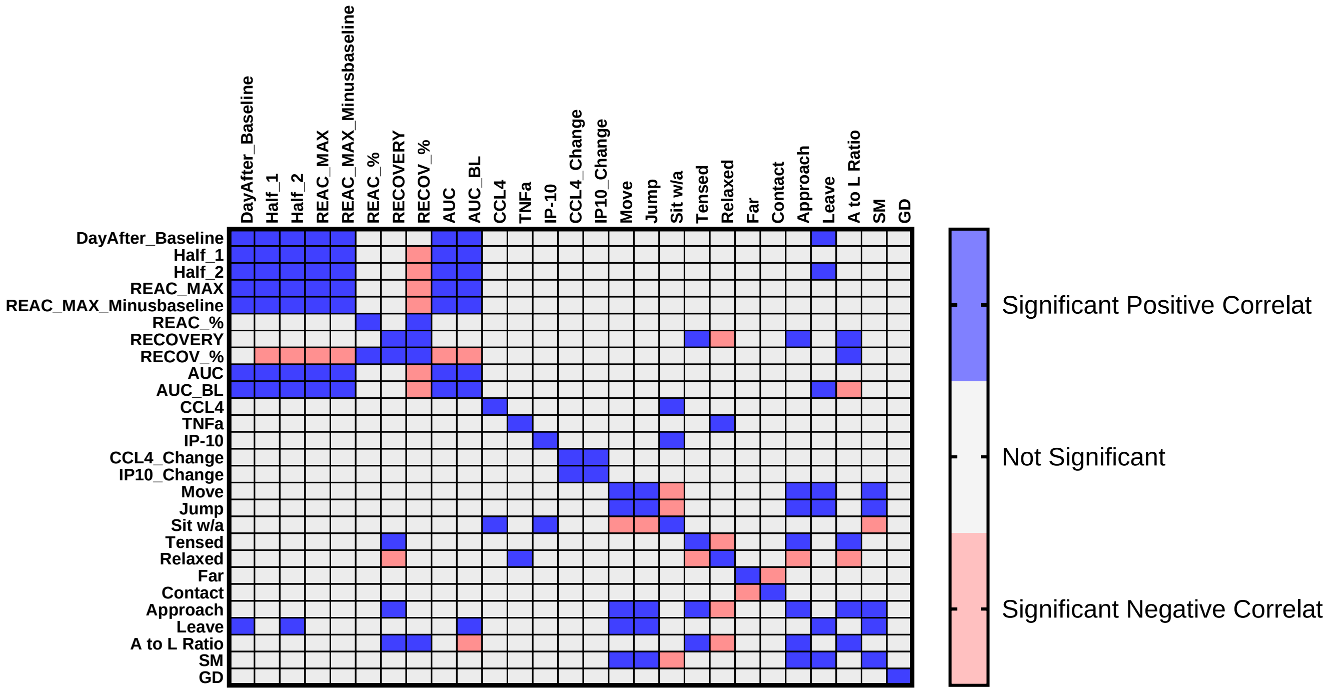


**Figure S2**. Correlation heat map for representative HPA and social reunion parameters across all 10 SSCs.

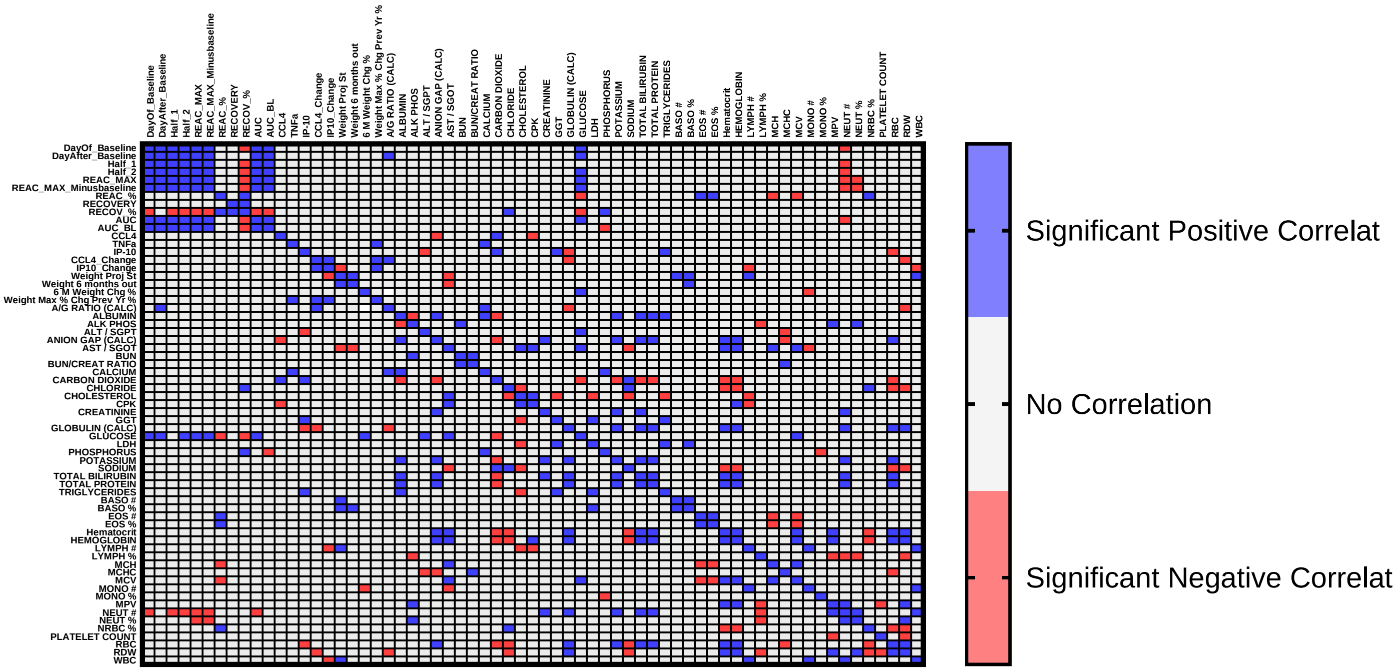


**Figure S3**. Correlation heat map for HPA and blood CBC/chemistry parameters.

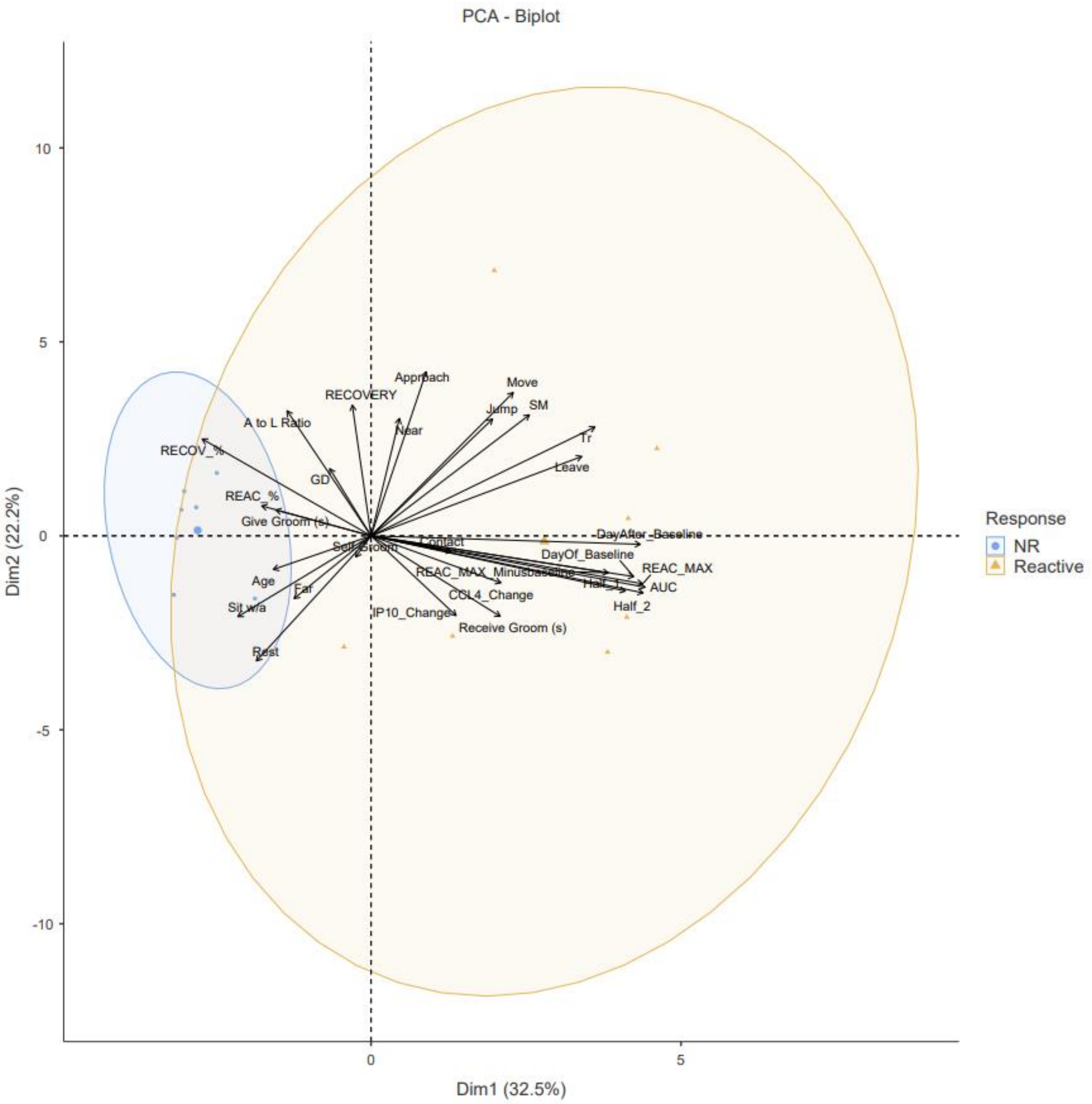


**Figure S4**. PCA biplot figure of social reunion behavior and cytokine data grouped by individuals who were reactor (orange) and non-reactor (blue) in their HPA responses.

Independent Samples T-Test by Age

| | | Statistic | df | p | | Effect Size |
|---|---|---|---|---|---|---|
| Pair_Length | Student's t | -0.81236 | 12.0 | 0.432 | Cohen's d | -0.43423 |
| Previous Partners (PP) | Student's t | -0.34133 | 12.0 | 0.739 | Cohen's d | -0.18245 |
| DayOf_Baseline | Student's t | 0.44711 | 12.0 | 0.663 | Cohen's d | 0.23899 |
| 8 | Student's t | 0.22736 | 12.0 | 0.824 | Cohen's d | 0.12153 |
| 9 | Student's t | 0.62481 | 12.0 | 0.544 | Cohen's d | 0.33397 |
| 10 | Student's t | 0.79424 | 12.0 | 0.442 | Cohen's d | 0.42454 |
| 11 | Student's t | 0.73962 | 12.0 | 0.474 | Cohen's d | 0.39535 |
| DayAfter_Baseline | Student's t | 0.38659 | 12.0 | 0.706 | Cohen's d | 0.20664 |
| Half_1 | Student's t | 0.39127 | 12.0 | 0.702 | Cohen's d | 0.20914 |
| Half_2 | Student's t | 0.80743 | 12.0 | 0.435 | Cohen's d | 0.43159 |
| REAC_MAX | Student's t | 0.62759 | 12.0 | 0.542 | Cohen's d | 0.33546 |
| REAC_MAX_Minusbaseline | Student's t | 0.71430 | 12.0 | 0.489 | Cohen's d | 0.38181 |
| REAC_% | Student's t | 0.39282 | 12.0 | 0.701 | Cohen's d | 0.20997 |
| CORT_ACCUM | Student's t | 0.64896 | 12.0 | 0.529 | Cohen's d | 0.34689 |
| RECOVERY | Student's t | -0.20221 | 12.0 | 0.843 | Cohen's d | -0.10808 |
| RECOV_% | Student's t | -0.01495 | 12.0 | 0.988 | Cohen's d | -0.00799 |
| AUC | Student's t | 0.61205 | 12.0 | 0.552 | Cohen's d | 0.32716 |
| AUC_BL | Student's t | 0.82967 | 12.0 | 0.423 | Cohen's d | 0.44348 |
| CCL4 | Student's t | -0.66749 | 12.0 | 0.517 | Cohen's d | -0.35679 |
| TNFa | Student's t | 0.65542[a] | 12.0 | 0.525 | Cohen's d | 0.35034 |
| IP-10 | Student's t | 0.34208 | 12.0 | 0.738 | Cohen's d | 0.18285 |
| CCL4_Change | Student's t | 1.82305 | 12.0 | 0.093 | Cohen's d | 0.97446 |
| IP10_Change | Student's t | 1.87440 | 12.0 | 0.085 | Cohen's d | 1.00191 |
| Rest | Student's t | -0.22566 | 12.0 | 0.825 | Cohen's d | -0.12062 |
| Move | Student's t | 0.52827[a] | 12.0 | 0.607 | Cohen's d | 0.28237 |
| Eat | Student's t | -0.70829 | 12.0 | 0.492 | Cohen's d | -0.37860 |
| Jump | Student's t | 1.06364[a] | 12.0 | 0.308 | Cohen's d | 0.56854 |
| Hang | Student's t | -0.51628 | 12.0 | 0.615 | Cohen's d | -0.27596 |
| Sit w/o | Student's t | 1.27499[a] | 12.0 | 0.226 | Cohen's d | 0.68151 |
| Sit w/a | Student's t | -0.16004 | 12.0 | 0.876 | Cohen's d | -0.08554 |
| 1 | Student's t | -0.91408 | 12.0 | 0.379 | Cohen's d | -0.48859 |
| 2 | Student's t | -1.31127 | 12.0 | 0.214 | Cohen's d | -0.70090 |
| 3 | Student's t | 1.66918 | 12.0 | 0.121 | Cohen's d | 0.89222 |
| 4 | Student's t | 0.67937 | 12.0 | 0.510 | Cohen's d | 0.36314 |
| Tr | Student's t | 0.34029 | 12.0 | 0.740 | Cohen's d | 0.18189 |
| Tensed | Student's t | -0.61863[a] | 11.0 | 0.549 | Cohen's d | -0.34417 |
| Relaxed | Student's t | 0.48671 | 11.0 | 0.636 | Cohen's d | 0.27078 |
| OV | Student's t | 0.53100 | 11.0 | 0.606 | Cohen's d | 0.29542 |
| Far | Student's t | -1.61475 | 12.0 | 0.132 | Cohen's d | -0.86312 |

Independent Samples T-Test by Age

| | | Statistic | df | p | | Effect Size |
|---|---|---|---|---|---|---|
| Near | Student's t | 0.81256 | 12.0 | 0.432 | Cohen's d | 0.43433 |
| Contact | Student's t | 1.47111 | 12.0 | 0.167 | Cohen's d | 0.78634 |
| Approach | Student's t | 0.46389 | 12.0 | 0.651 | Cohen's d | 0.24796 |
| Approach latency (s) | Student's t | 0.12708 | 12.0 | 0.901 | Cohen's d | 0.06793 |
| Leave | Student's t | 0.78495 | 12.0 | 0.448 | Cohen's d | 0.41957 |
| Leave Latency (s) | Student's t | -0.69801 | 12.0 | 0.498 | Cohen's d | -0.37310 |
| A to L Ratio | Student's t | -0.54436 | 12.0 | 0.596 | Cohen's d | -0.29098 |
| Receive Groom | Student's t | 0.68192 | 12.0 | 0.508 | Cohen's d | 0.36450 |
| Receive Groom (s) | Student's t | 0.24892 | 12.0 | 0.808 | Cohen's d | 0.13306 |
| Give Groom | Student's t | 1.06066 | 12.0 | 0.310 | Cohen's d | 0.56695 |
| Give Groom (s) | Student's t | 0.85993 | 12.0 | 0.407 | Cohen's d | 0.45965 |
| Mate | Student's t | -0.61803 | 12.0 | 0.548 | Cohen's d | -0.33035 |
| Self-Groom | Student's t | -0.17486 | 12.0 | 0.864 | Cohen's d | -0.09347 |
| SM | Student's t | -0.05077 | 12.0 | 0.960 | Cohen's d | -0.02714 |
| GD | Student's t | 0.90527 | 12.0 | 0.383 | Cohen's d | 0.48389 |
| Rest (2) | Student's t | 1.30766 | 12.0 | 0.215 | Cohen's d | 0.69898 |
| Move (2) | Student's t | -1.11554 | 12.0 | 0.286 | Cohen's d | -0.59628 |
| Eat (2) | Student's t | -1.45056 | 12.0 | 0.173 | Cohen's d | -0.77536 |
| Jump (2) | Student's t | -0.78077 | 12.0 | 0.450 | Cohen's d | -0.41734 |
| Sit | Student's t | -0.00308 | 12.0 | 0.998 | Cohen's d | -0.00165 |
| Stand | Student's t | 0.18853 | 12.0 | 0.854 | Cohen's d | 0.10077 |
| Tr (2) | Student's t | -0.09861 | 12.0 | 0.923 | Cohen's d | -0.05271 |
| Tensed (2) | Student's t | 0.17289 | 11.0 | 0.866 | Cohen's d | 0.09618 |
| Relaxed (2) | Student's t | -0.14832 | 11.0 | 0.885 | Cohen's d | -0.08252 |
| Head Scans | Student's t | 1.98366 | 12.0 | 0.071 | Cohen's d | 1.06031 |
| SM (2) | Student's t | -1.29345 | 12.0 | 0.220 | Cohen's d | -0.69138 |
| Self Groom | Student's t | -0.33410 | 12.0 | 0.744 | Cohen's d | -0.17858 |
| In Bucket | Student's t | 0.57095 | 12.0 | 0.579 | Cohen's d | 0.30519 |
| Shrill | Student's t | 0.43881 | 12.0 | 0.669 | Cohen's d | 0.23455 |
| BCS | Student's t | 0.37354 | 12.0 | 0.715 | Cohen's d | 0.19967 |
| Weight | Student's t | 0.53485 | 12.0 | 0.603 | Cohen's d | 0.28589 |
| Weight Proj St | Student's t | -0.09713 | 12.0 | 0.924 | Cohen's d | -0.05192 |
| Weight 6 months out | Student's t | 0.93262 | 12.0 | 0.369 | Cohen's d | 0.49851 |
| 6 M Weight Chg % | Student's t | 1.81582 | 12.0 | 0.094 | Cohen's d | 0.97060 |
| Weight Max % Chg Prev Yr % | Student's t | 2.86371 | 12.0 | 0.014 | Cohen's d | 1.53072 |
| A/G RATIO (CALC) | Student's t | 0.31634 | 12.0 | 0.757 | Cohen's d | 0.16909 |
| ALBUMIN | Student's t | 0.44620 | 12.0 | 0.663 | Cohen's d | 0.23851 |
| ALK PHOS | Student's t | -0.29807 | 12.0 | 0.771 | Cohen's d | -0.15933 |
| ALT / SGPT | Student's t | -0.11881 | 12.0 | 0.907 | Cohen's d | -0.06351 |
| ANION GAP (CALC) | Student's t | -0.38007 | 12.0 | 0.711 | Cohen's d | -0.20315 |
| AST / SGOT | Student's t | 0.04127 | 12.0 | 0.968 | Cohen's d | 0.02206 |

Independent Samples T-Test by Age

| | | Statistic | df | p | | Effect Size |
|---|---|---|---|---|---|---|
| BUN | Student's t | -0.70605 | 12.0 | 0.494 | Cohen's d | -0.37740 |
| BUN/CREAT RATIO | Student's t | 0.74477 | 12.0 | 0.471 | Cohen's d | 0.39809 |
| CALCIUM | Student's t | -0.06110 | 12.0 | 0.952 | Cohen's d | -0.03266 |
| CARBON DIOXIDE | Student's t | 1.51414 | 12.0 | 0.156 | Cohen's d | 0.80934 |
| CHLORIDE | Student's t | -0.89893 | 12.0 | 0.386 | Cohen's d | -0.48050 |
| CHOLESTEROL | Student's t | -0.67452 | 12.0 | 0.513 | Cohen's d | -0.36055 |
| CPK | Student's t | -0.45072 | 11.0 | 0.661 | Cohen's d | -0.25076 |
| CREATININE | Student's t | -0.75574 | 12.0 | 0.464 | Cohen's d | -0.40396 |
| GGT | Student's t | -1.75487[a] | 11.0 | 0.107 | Cohen's d | -0.97632 |
| GLOBULIN (CALC) | Student's t | 0.18162 | 12.0 | 0.859 | Cohen's d | 0.09708 |
| GLUCOSE | Student's t | 0.81306 | 12.0 | 0.432 | Cohen's d | 0.43460 |
| LDH | Student's t | -0.17062 | 11.0 | 0.868 | Cohen's d | -0.09492 |
| PHOSPHORUS | Student's t | -0.29605 | 12.0 | 0.772 | Cohen's d | -0.15824 |
| POTASSIUM | Student's t | 1.19811 | 12.0 | 0.254 | Cohen's d | 0.64042 |
| SODIUM | Student's t | -0.31506 | 12.0 | 0.758 | Cohen's d | -0.16841 |
| TOTAL BILIRUBIN | Student's t | 1.27920[a] | 12.0 | 0.225 | Cohen's d | 0.68376 |
| TOTAL PROTEIN | Student's t | 0.40670 | 12.0 | 0.691 | Cohen's d | 0.21739 |
| TRIGLYCERIDES | Student's t | 0.06284 | 11.0 | 0.951 | Cohen's d | 0.03496 |
| BASO # | Student's t | 0.61237 | 12.0 | 0.552 | Cohen's d | 0.32733 |
| BASO % | Student's t | 1.15214 | 12.0 | 0.272 | Cohen's d | 0.61585 |
| EOS # | Student's t | 1.47029 | 12.0 | 0.167 | Cohen's d | 0.78591 |
| EOS % | Student's t | 1.47625 | 12.0 | 0.166 | Cohen's d | 0.78909 |
| Hematocrit | Student's t | 1.51387 | 12.0 | 0.156 | Cohen's d | 0.80920 |
| HEMOGLOBIN | Student's t | 1.61155[a] | 12.0 | 0.133 | Cohen's d | 0.86141 |
| LYMPH # | Student's t | -0.58519 | 12.0 | 0.569 | Cohen's d | -0.31280 |
| LYMPH % | Student's t | -0.24082 | 12.0 | 0.814 | Cohen's d | -0.12872 |
| MCH | Student's t | -2.14394 | 12.0 | 0.053 | Cohen's d | -1.14599 |
| MCHC | Student's t | -0.22706 | 12.0 | 0.824 | Cohen's d | -0.12137 |
| MCV | Student's t | -1.49277 | 12.0 | 0.161 | Cohen's d | -0.79792 |
| MONO # | Student's t | -0.96225 | 12.0 | 0.355 | Cohen's d | -0.51434 |
| MONO % | Student's t | -0.72824 | 12.0 | 0.480 | Cohen's d | -0.38926 |
| MPV | Student's t | 0.04308 | 12.0 | 0.966 | Cohen's d | 0.02303 |
| NEUT # | Student's t | -0.43697[a] | 12.0 | 0.670 | Cohen's d | -0.23357 |
| NEUT % | Student's t | 0.17079 | 12.0 | 0.867 | Cohen's d | 0.09129 |
| NRBC % | Student's t | -0.07981 | 12.0 | 0.938 | Cohen's d | -0.04266 |
| PLATELET COUNT | Student's t | 0.40738 | 12.0 | 0.691 | Cohen's d | 0.21775 |
| RBC | Student's t | 2.14882[a] | 12.0 | 0.053 | Cohen's d | 1.14859 |
| RDW | Student's t | 0.43823 | 12.0 | 0.669 | Cohen's d | 0.23424 |
| WBC | Student's t | -0.80715 | 12.0 | 0.435 | Cohen's d | -0.43144 |
| A/G RATIO (CALC) (2) | Student's t | -0.80937 | 11.0 | 0.435 | Cohen's d | -0.45029 |
| ALBUMIN (2) | Student's t | -0.63776 | 11.0 | 0.537 | Cohen's d | -0.35482 |

Independent Samples T-Test by Age

| | | Statistic | df | p | | Effect Size |
|---|---|---|---|---|---|---|
| ALK PHOS (2) | Student's t | 0.44809 | 11.0 | 0.663 | Cohen's d | 0.24930 |
| ALT / SGPT (2) | Student's t | -0.66667 | 11.0 | 0.519 | Cohen's d | -0.37090 |
| ANION GAP (CALC) (2) | Student's t | -0.22253 | 11.0 | 0.828 | Cohen's d | -0.12380 |
| AST / SGOT (2) | Student's t | -0.30189 | 11.0 | 0.768 | Cohen's d | -0.16796 |
| BUN (2) | Student's t | -0.43192 | 11.0 | 0.674 | Cohen's d | -0.24030 |
| BUN/CREAT RATIO (2) | Student's t | -0.10406 | 11.0 | 0.919 | Cohen's d | -0.05789 |
| CALCIUM (2) | Student's t | -0.80204 | 11.0 | 0.440 | Cohen's d | -0.44621 |
| CARBON DIOXIDE (2) | Student's t | -0.10519 | 11.0 | 0.918 | Cohen's d | -0.05852 |
| CHLORIDE (2) | Student's t | -0.96976 | 11.0 | 0.353 | Cohen's d | -0.53953 |
| CHOLESTEROL (2) | Student's t | 1.30499[a] | 11.0 | 0.219 | Cohen's d | 0.72603 |
| CPK (2) | Student's t | 0.48322 | 11.0 | 0.638 | Cohen's d | 0.26884 |
| CREATININE (2) | Student's t | -1.03304 | 11.0 | 0.324 | Cohen's d | -0.57473 |
| GGT (2) | Student's t | -0.26842 | 11.0 | 0.793 | Cohen's d | -0.14933 |
| GLOBULIN (CALC) (2) | Student's t | 0.02753 | 11.0 | 0.979 | Cohen's d | 0.01532 |
| GLUCOSE (2) | Student's t | 0.42439 | 11.0 | 0.679 | Cohen's d | 0.23611 |
| LDH (2) | Student's t | -1.17352 | 11.0 | 0.265 | Cohen's d | -0.65289 |
| PHOSPHORUS (2) | Student's t | -0.84268 | 11.0 | 0.417 | Cohen's d | -0.46882 |
| POTASSIUM (2) | Student's t | -0.28647 | 11.0 | 0.780 | Cohen's d | -0.15938 |
| SODIUM (2) | Student's t | -1.20460 | 11.0 | 0.254 | Cohen's d | -0.67018 |
| TOTAL BILIRUBIN (2) | Student's t | 0.02800 | 11.0 | 0.978 | Cohen's d | 0.01558 |
| TOTAL PROTEIN (2) | Student's t | -0.38801 | 11.0 | 0.705 | Cohen's d | -0.21587 |
| TRIGLYCERIDES (2) | Student's t | 0.24848 | 11.0 | 0.808 | Cohen's d | 0.13824 |
| BASO # (2) | Student's t | -0.59027 | 11.0 | 0.567 | Cohen's d | -0.32839 |
| BASO % (2) | Student's t | -0.44733 | 11.0 | 0.663 | Cohen's d | -0.24887 |
| EOS # (2) | Student's t | 2.02988 | 11.0 | 0.067 | Cohen's d | 1.12932 |
| EOS % (2) | Student's t | 1.60414 | 11.0 | 0.137 | Cohen's d | 0.89246 |
| Hematocrit (2) | Student's t | -0.04870 | 11.0 | 0.962 | Cohen's d | -0.02709 |
| HEMOGLOBIN (2) | Student's t | -0.02669 | 11.0 | 0.979 | Cohen's d | -0.01485 |
| LYMPH # (2) | Student's t | -2.04110[a] | 11.0 | 0.066 | Cohen's d | -1.13556 |
| LYMPH % (2) | Student's t | -1.34947[a] | 11.0 | 0.204 | Cohen's d | -0.75078 |
| MCH (2) | Student's t | -0.49388 | 11.0 | 0.631 | Cohen's d | -0.27477 |
| MCHC (2) | Student's t | -0.02679 | 11.0 | 0.979 | Cohen's d | -0.01491 |
| MCV (2) | Student's t | -0.41753 | 11.0 | 0.684 | Cohen's d | -0.23229 |
| MONO # (2) | Student's t | -0.77352[a] | 11.0 | 0.456 | Cohen's d | -0.43034 |
| MONO % (2) | Student's t | 0.53977 | 11.0 | 0.600 | Cohen's d | 0.30030 |
| MPV (2) | Student's t | -0.45097 | 11.0 | 0.661 | Cohen's d | -0.25089 |
| NEUT # (2) | Student's t | -0.13005 | 11.0 | 0.899 | Cohen's d | -0.07235 |
| NEUT % (2) | Student's t | 0.35668 | 11.0 | 0.728 | Cohen's d | 0.19844 |
| NRBC % (2) | Student's t | 1.39749 | 11.0 | 0.190 | Cohen's d | 0.77749 |
| PLATELET COUNT (2) | Student's t | 1.00934 | 11.0 | 0.334 | Cohen's d | 0.56155 |
| RBC (2) | Student's t | 0.18013 | 11.0 | 0.860 | Cohen's d | 0.10022 |

Independent Samples T-Test by Age

| | | Statistic | df | p | | Effect Size |
|---|---|---|---|---|---|---|
| RDW (2) | Student's t | -0.04123 | 11.0 | 0.968 | Cohen's d | -0.02294 |
| WBC (2) | Student's t | -1.48989[a] | 11.0 | 0.164 | Cohen's d | -0.82890 |

Note. $H_a$ $\mu_{Peri\text{-}Geri} \neq \mu_{Very\text{-}Geri}$

[a] Levene's test is significant ($p < .05$), suggesting a violation of the assumption of equal variances

**SI Table 1.** T-Tests for each measured parameter based on age group. Behavioral variables followed by “(2)” are isolation behaviors. Blood CBC/Chem parameters are averaged or parameters followed by “(2)” are change from post-pre. Positive t-statistics indicate peri-geri > very-geri. For clarification on Behavioral variable names, see Table 1.

Independent Samples T-Test by Sex

| | | Statistic | df | p | | Effect Size |
|---|---|---|---|---|---|---|
| Pair_Length | Student's t | -1.6867 | 12.0 | 0.117 | Cohen's d | -0.91094 |
| Previous Partners (PP) | Student's t | 0.6484 | 12.0 | 0.529 | Cohen's d | 0.35020 |
| DayOf_Baseline | Student's t | 0.8653 | 12.0 | 0.404 | Cohen's d | 0.46734 |
| 8 | Student's t | 0.4576 | 12.0 | 0.655 | Cohen's d | 0.24715 |
| 9 | Student's t | 1.0659 | 12.0 | 0.307 | Cohen's d | 0.57563 |
| 10 | Student's t | 1.4327 | 12.0 | 0.177 | Cohen's d | 0.77374 |
| 11 | Student's t | 1.6382 | 12.0 | 0.127 | Cohen's d | 0.88475 |
| DayAfter_Baseline | Student's t | 0.6210 | 12.0 | 0.546 | Cohen's d | 0.33536 |
| Half_1 | Student's t | 0.7342 | 12.0 | 0.477 | Cohen's d | 0.39650 |
| Half_2 | Student's t | 1.5320 | 12.0 | 0.151 | Cohen's d | 0.82738 |
| REAC_MAX | Student's t | 1.1906 | 12.0 | 0.257 | Cohen's d | 0.64298 |
| REAC_MAX_Minusbaseline | Student's t | 1.1728 | 12.0 | 0.264 | Cohen's d | 0.63340 |
| REAC_% | Student's t | -0.5674 | 12.0 | 0.581 | Cohen's d | -0.30645 |
| CORT_ACCUM | Student's t | 1.1828 | 12.0 | 0.260 | Cohen's d | 0.63879 |
| RECOVERY | Student's t | -1.7496[a] | 12.0 | 0.106 | Cohen's d | -0.94489 |
| RECOV_% | Student's t | -2.2466 | 12.0 | 0.044 | Cohen's d | -1.21329 |
| AUC | Student's t | 1.1603 | 12.0 | 0.268 | Cohen's d | 0.62662 |
| AUC_BL | Student's t | 1.6508 | 12.0 | 0.125 | Cohen's d | 0.89152 |
| CCL4 | Student's t | -0.0556[a] | 12.0 | 0.957 | Cohen's d | -0.03003 |
| TNFa | Student's t | 0.0432 | 12.0 | 0.966 | Cohen's d | 0.02333 |
| IP-10 | Student's t | -0.9526 | 12.0 | 0.360 | Cohen's d | -0.51448 |
| CCL4_Change | Student's t | 0.2293 | 12.0 | 0.822 | Cohen's d | 0.12386 |
| IP10_Change | Student's t | 0.1019 | 12.0 | 0.920 | Cohen's d | 0.05505 |
| Rest | Student's t | -0.1029 | 12.0 | 0.920 | Cohen's d | -0.05556 |
| Move | Student's t | -0.3096 | 12.0 | 0.762 | Cohen's d | -0.16720 |
| Eat | Student's t | 0.9641 | 12.0 | 0.354 | Cohen's d | 0.52065 |
| Jump | Student's t | -0.6598 | 12.0 | 0.522 | Cohen's d | -0.35634 |
| Hang | Student's t | -0.0885 | 12.0 | 0.931 | Cohen's d | -0.04778 |
| Sit w/o | Student's t | 0.7765 | 12.0 | 0.453 | Cohen's d | 0.41934 |
| Sit w/a | Student's t | 0.4219 | 12.0 | 0.681 | Cohen's d | 0.22783 |
| 1 | Student's t | 0.5756 | 12.0 | 0.576 | Cohen's d | 0.31084 |
| 2 | Student's t | -1.3383 | 12.0 | 0.206 | Cohen's d | -0.72276 |
| 3 | Student's t | 0.7398 | 12.0 | 0.474 | Cohen's d | 0.39953 |
| 4 | Student's t | -0.0385[a] | 12.0 | 0.970 | Cohen's d | -0.02079 |
| Tr | Student's t | -0.1412 | 12.0 | 0.890 | Cohen's d | -0.07623 |
| Tensed | Student's t | -1.3488 | 11.0 | 0.205 | Cohen's d | -0.75039 |
| Relaxed | Student's t | 1.2031 | 11.0 | 0.254 | Cohen's d | 0.66935 |
| OV | Student's t | 0.7120 | 11.0 | 0.491 | Cohen's d | 0.39613 |
| Far | Student's t | 0.7924 | 12.0 | 0.444 | Cohen's d | 0.42792 |
| Near | Student's t | -1.4584 | 12.0 | 0.170 | Cohen's d | -0.78764 |

Independent Samples T-Test by Sex

| | | Statistic | df | p | | Effect Size |
|---|---|---|---|---|---|---|
| Contact | Student's t | 0.0871 | 12.0 | 0.932 | Cohen's d | 0.04704 |
| Approach | Student's t | -2.2652 | 12.0 | 0.043 | Cohen's d | -1.22333 |
| Approach latency (s) | Student's t | 4.5850[a] | 12.0 | <.001 | Cohen's d | 2.47616 |
| Leave | Student's t | 0.7614 | 12.0 | 0.461 | Cohen's d | 0.41118 |
| Leave Latency (s) | Student's t | -1.3906 | 12.0 | 0.190 | Cohen's d | -0.75103 |
| A to L Ratio | Student's t | -5.0360[a] | 12.0 | <.001 | Cohen's d | -2.71975 |
| Receive Groom | Student's t | 0.9514 | 12.0 | 0.360 | Cohen's d | 0.51383 |
| Receive Groom (s) | Student's t | 0.5779 | 12.0 | 0.574 | Cohen's d | 0.31207 |
| Give Groom | Student's t | -0.6968 | 12.0 | 0.499 | Cohen's d | -0.37632 |
| Give Groom (s) | Student's t | -0.7094 | 12.0 | 0.492 | Cohen's d | -0.38312 |
| Mate | Student's t | -1.9478[a] | 12.0 | 0.075 | Cohen's d | -1.05194 |
| Self-Groom | Student's t | 0.9216[a] | 12.0 | 0.375 | Cohen's d | 0.49775 |
| SM | Student's t | 0.1660 | 12.0 | 0.871 | Cohen's d | 0.08967 |
| GD | Student's t | -1.8771[a] | 12.0 | 0.085 | Cohen's d | -1.01376 |
| Rest (2) | Student's t | -1.2615 | 12.0 | 0.231 | Cohen's d | -0.68131 |
| Move (2) | Student's t | 1.7907 | 12.0 | 0.099 | Cohen's d | 0.96710 |
| Eat (2) | Student's t | -1.9981 | 12.0 | 0.069 | Cohen's d | -1.07909 |
| Jump (2) | Student's t | 2.2383 | 12.0 | 0.045 | Cohen's d | 1.20884 |
| Sit | Student's t | -2.8771 | 12.0 | 0.014 | Cohen's d | -1.55383 |
| Stand | Student's t | 2.6562 | 12.0 | 0.021 | Cohen's d | 1.43451 |
| Tr (2) | Student's t | 2.7315[a] | 12.0 | 0.018 | Cohen's d | 1.47517 |
| Tensed (2) | Student's t | 1.9577 | 11.0 | 0.076 | Cohen's d | 1.08915 |
| Relaxed (2) | Student's t | -1.9504 | 11.0 | 0.077 | Cohen's d | -1.08511 |
| Head Scans | Student's t | 0.9377 | 12.0 | 0.367 | Cohen's d | 0.50643 |
| SM (2) | Student's t | 2.0154[a] | 12.0 | 0.067 | Cohen's d | 1.08844 |
| Self Groom | Student's t | -0.4575[a] | 12.0 | 0.655 | Cohen's d | -0.24709 |
| In Bucket | Student's t | -0.1861 | 12.0 | 0.855 | Cohen's d | -0.10052 |
| Shrill | Student's t | 0.4383 | 12.0 | 0.669 | Cohen's d | 0.23672 |
| BCS | Student's t | 0.1073 | 12.0 | 0.916 | Cohen's d | 0.05793 |
| Weight | Student's t | 0.5605 | 12.0 | 0.585 | Cohen's d | 0.30271 |
| Weight Proj St | Student's t | 0.4852 | 12.0 | 0.636 | Cohen's d | 0.26203 |
| Weight 6 months out | Student's t | 1.3841 | 12.0 | 0.192 | Cohen's d | 0.74751 |
| 6 M Weight Chg % | Student's t | 1.4940 | 12.0 | 0.161 | Cohen's d | 0.80686 |
| Weight Max % Chg Prev Yr % | Student's t | 0.1671 | 12.0 | 0.870 | Cohen's d | 0.09024 |
| A/G RATIO (CALC) | Student's t | -0.5391 | 12.0 | 0.600 | Cohen's d | -0.29112 |
| ALBUMIN | Student's t | -0.3495 | 12.0 | 0.733 | Cohen's d | -0.18877 |
| ALK PHOS | Student's t | 0.2808 | 12.0 | 0.784 | Cohen's d | 0.15168 |
| ALT / SGPT | Student's t | 1.6779 | 12.0 | 0.119 | Cohen's d | 0.90615 |
| ANION GAP (CALC) | Student's t | 0.5505 | 12.0 | 0.592 | Cohen's d | 0.29731 |
| AST / SGOT | Student's t | -1.0082 | 12.0 | 0.333 | Cohen's d | -0.54448 |
| BUN | Student's t | 0.9677 | 12.0 | 0.352 | Cohen's d | 0.52260 |

Independent Samples T-Test by Sex

| | | Statistic | df | p | | Effect Size |
|---|---|---|---|---|---|---|
| BUN/CREAT RATIO | Student's t | 0.6890 | 12.0 | 0.504 | Cohen's d | 0.37210 |
| CALCIUM | Student's t | -0.0764 | 12.0 | 0.940 | Cohen's d | -0.04128 |
| CARBON DIOXIDE | Student's t | -0.7448 | 12.0 | 0.471 | Cohen's d | -0.40226 |
| CHLORIDE | Student's t | -0.4689 | 12.0 | 0.648 | Cohen's d | -0.25325 |
| CHOLESTEROL | Student's t | -1.6075 | 12.0 | 0.134 | Cohen's d | -0.86815 |
| CPK | Student's t | -1.0423[a] | 11.0 | 0.320 | Cohen's d | -0.59419 |
| CREATININE | Student's t | 0.5588 | 12.0 | 0.587 | Cohen's d | 0.30179 |
| GGT | Student's t | -0.4506 | 11.0 | 0.661 | Cohen's d | -0.25690 |
| GLOBULIN (CALC) | Student's t | 0.4309 | 12.0 | 0.674 | Cohen's d | 0.23271 |
| GLUCOSE | Student's t | 0.4723 | 12.0 | 0.645 | Cohen's d | 0.25507 |
| LDH | Student's t | -0.5282 | 11.0 | 0.608 | Cohen's d | -0.30111 |
| PHOSPHORUS | Student's t | 0.8187 | 12.0 | 0.429 | Cohen's d | 0.44216 |
| POTASSIUM | Student's t | 0.1368 | 12.0 | 0.893 | Cohen's d | 0.07388 |
| SODIUM | Student's t | -0.4878 | 12.0 | 0.634 | Cohen's d | -0.26347 |
| TOTAL BILIRUBIN | Student's t | -0.2315 | 12.0 | 0.821 | Cohen's d | -0.12500 |
| TOTAL PROTEIN | Student's t | 0.0000 | 12.0 | 1.000 | Cohen's d | 0.00000 |
| TRIGLYCERIDES | Student's t | 0.0717[a] | 11.0 | 0.944 | Cohen's d | 0.04087 |
| BASO # | Student's t | 0.3499 | 12.0 | 0.732 | Cohen's d | 0.18898 |
| BASO % | Student's t | -1.1377 | 12.0 | 0.277 | Cohen's d | -0.61444 |
| EOS # | Student's t | 0.1676 | 12.0 | 0.870 | Cohen's d | 0.09054 |
| EOS % | Student's t | 0.2322 | 12.0 | 0.820 | Cohen's d | 0.12539 |
| Hematocrit | Student's t | -1.5145 | 12.0 | 0.156 | Cohen's d | -0.81794 |
| HEMOGLOBIN | Student's t | -1.5287 | 12.0 | 0.152 | Cohen's d | -0.82562 |
| LYMPH # | Student's t | 1.1266 | 12.0 | 0.282 | Cohen's d | 0.60845 |
| LYMPH % | Student's t | 1.6891[a] | 12.0 | 0.117 | Cohen's d | 0.91220 |
| MCH | Student's t | 0.8168 | 12.0 | 0.430 | Cohen's d | 0.44112 |
| MCHC | Student's t | 0.6453[a] | 12.0 | 0.531 | Cohen's d | 0.34853 |
| MCV | Student's t | 0.3999 | 12.0 | 0.696 | Cohen's d | 0.21595 |
| MONO # | Student's t | 1.3833 | 12.0 | 0.192 | Cohen's d | 0.74709 |
| MONO % | Student's t | 0.6386 | 12.0 | 0.535 | Cohen's d | 0.34488 |
| MPV | Student's t | -0.1494 | 12.0 | 0.884 | Cohen's d | -0.08067 |
| NEUT # | Student's t | -1.3165 | 12.0 | 0.213 | Cohen's d | -0.71100 |
| NEUT % | Student's t | -1.7668[a] | 12.0 | 0.103 | Cohen's d | -0.95421 |
| NRBC % | Student's t | 0.3457 | 12.0 | 0.736 | Cohen's d | 0.18669 |
| PLATELET COUNT | Student's t | -0.6446 | 12.0 | 0.531 | Cohen's d | -0.34813 |
| RBC | Student's t | -1.7299 | 12.0 | 0.109 | Cohen's d | -0.93425 |
| RDW | Student's t | 0.9241 | 12.0 | 0.374 | Cohen's d | 0.49908 |
| WBC | Student's t | 0.6943 | 12.0 | 0.501 | Cohen's d | 0.37497 |
| A/G RATIO (CALC) (2) | Student's t | -0.0168 | 11.0 | 0.987 | Cohen's d | -0.00957 |
| ALBUMIN (2) | Student's t | -1.6195 | 11.0 | 0.134 | Cohen's d | -0.92323 |
| ALK PHOS (2) | Student's t | 1.3496 | 11.0 | 0.204 | Cohen's d | 0.76938 |

Independent Samples T-Test by Sex

| | | Statistic | df | p | | Effect Size |
|---|---|---|---|---|---|---|
| ALT / SGPT (2) | Student's t | 1.1255 | 11.0 | 0.284 | Cohen's d | 0.64164 |
| ANION GAP (CALC) (2) | Student's t | -0.5287 | 11.0 | 0.608 | Cohen's d | -0.30141 |
| AST / SGOT (2) | Student's t | 0.9820 | 11.0 | 0.347 | Cohen's d | 0.55982 |
| BUN (2) | Student's t | 0.8134[a] | 11.0 | 0.433 | Cohen's d | 0.46371 |
| BUN/CREAT RATIO (2) | Student's t | 0.5220[a] | 11.0 | 0.612 | Cohen's d | 0.29760 |
| CALCIUM (2) | Student's t | -0.3825 | 11.0 | 0.709 | Cohen's d | -0.21808 |
| CARBON DIOXIDE (2) | Student's t | 0.4346 | 11.0 | 0.672 | Cohen's d | 0.24778 |
| CHLORIDE (2) | Student's t | -0.9621 | 11.0 | 0.357 | Cohen's d | -0.54846 |
| CHOLESTEROL (2) | Student's t | 1.3706[a] | 11.0 | 0.198 | Cohen's d | 0.78136 |
| CPK (2) | Student's t | -0.2613 | 11.0 | 0.799 | Cohen's d | -0.14896 |
| CREATININE (2) | Student's t | 0.0000 | 11.0 | 1.000 | Cohen's d | 0.00000 |
| GGT (2) | Student's t | -1.2322 | 11.0 | 0.244 | Cohen's d | -0.70247 |
| GLOBULIN (CALC) (2) | Student's t | -0.9276 | 11.0 | 0.374 | Cohen's d | -0.52882 |
| GLUCOSE (2) | Student's t | 1.7912 | 11.0 | 0.101 | Cohen's d | 1.02112 |
| LDH (2) | Student's t | -0.9284 | 11.0 | 0.373 | Cohen's d | -0.52926 |
| PHOSPHORUS (2) | Student's t | -1.2794 | 11.0 | 0.227 | Cohen's d | -0.72938 |
| POTASSIUM (2) | Student's t | -1.8074 | 11.0 | 0.098 | Cohen's d | -1.03038 |
| SODIUM (2) | Student's t | -0.6124 | 11.0 | 0.553 | Cohen's d | -0.34913 |
| TOTAL BILIRUBIN (2) | Student's t | -1.0871 | 11.0 | 0.300 | Cohen's d | -0.61972 |
| TOTAL PROTEIN (2) | Student's t | -1.6631 | 11.0 | 0.124 | Cohen's d | -0.94814 |
| TRIGLYCERIDES (2) | Student's t | -1.5213 | 11.0 | 0.156 | Cohen's d | -0.86727 |
| BASO # (2) | Student's t | 1.7262 | 11.0 | 0.112 | Cohen's d | 0.98411 |
| BASO % (2) | Student's t | 0.7659 | 11.0 | 0.460 | Cohen's d | 0.43665 |
| EOS # (2) | Student's t | -1.9470 | 11.0 | 0.078 | Cohen's d | -1.10996 |
| EOS % (2) | Student's t | -1.5870 | 11.0 | 0.141 | Cohen's d | -0.90473 |
| Hematocrit (2) | Student's t | 0.1387 | 11.0 | 0.892 | Cohen's d | 0.07909 |
| HEMOGLOBIN (2) | Student's t | 0.1155 | 11.0 | 0.910 | Cohen's d | 0.06587 |
| LYMPH # (2) | Student's t | -0.2757 | 11.0 | 0.788 | Cohen's d | -0.15715 |
| LYMPH % (2) | Student's t | 0.3378 | 11.0 | 0.742 | Cohen's d | 0.19256 |
| MCH (2) | Student's t | 1.0055 | 11.0 | 0.336 | Cohen's d | 0.57321 |
| MCHC (2) | Student's t | -0.1718 | 11.0 | 0.867 | Cohen's d | -0.09795 |
| MCV (2) | Student's t | 1.1828 | 11.0 | 0.262 | Cohen's d | 0.67429 |
| MONO # (2) | Student's t | -1.7443 | 11.0 | 0.109 | Cohen's d | -0.99440 |
| MONO % (2) | Student's t | -0.6044 | 11.0 | 0.558 | Cohen's d | -0.34456 |
| MPV (2) | Student's t | 1.2051 | 11.0 | 0.253 | Cohen's d | 0.68703 |
| NEUT # (2) | Student's t | -0.6176 | 11.0 | 0.549 | Cohen's d | -0.35210 |
| NEUT % (2) | Student's t | -0.6460 | 11.0 | 0.532 | Cohen's d | -0.36826 |
| NRBC % (2) | Student's t | -1.0070 | 11.0 | 0.336 | Cohen's d | -0.57409 |
| PLATELET COUNT (2) | Student's t | -1.5168 | 11.0 | 0.158 | Cohen's d | -0.86473 |
| RBC (2) | Student's t | -0.2464 | 11.0 | 0.810 | Cohen's d | -0.14048 |
| RDW (2) | Student's t | 0.2560 | 11.0 | 0.803 | Cohen's d | 0.14597 |

Independent Samples T-Test by Sex

| | | Statistic | df | p | | Effect Size |
|---|---|---|---|---|---|---|
| WBC (2) | Student's t | -0.6011 | 11.0 | 0.560 | Cohen's d | -0.34269 |

Note. $H_a$ $\mu_{Female} \neq \mu_{Male}$

[a] Levene's test is significant ($p < .05$), suggesting a violation of the assumption of equal variances

**SI Table 2.** T-Tests for each measured parameter based on sex. Behavioral variables followed by “(2)” are isolation behaviors. Blood CBC/Chem parameters are averaged or parameters followed by “(2)” are change from post-pre. Positive t-statistics indicate females > males. For clarification on Behavioral variable names, see Table 1.

Independent Samples T-Test by HPA Reactor

| | | Statistic | df | p | | Effect Size |
|---|---|---|---|---|---|---|
| Pair_Length | Student's t | -0.16426 | 12.0 | 0.872 | Cohen's d | -0.08780 |
| Previous Partners (PP) | Student's t | 0.34133 | 12.0 | 0.739 | Cohen's d | 0.18245 |
| DayOf_Baseline | Student's t | -4.34432[a] | 12.0 | <.001 | Cohen's d | -2.32213 |
| 8 | Student's t | -3.79642[a] | 12.0 | 0.003 | Cohen's d | -2.02927 |
| 9 | Student's t | -4.77571[a] | 12.0 | <.001 | Cohen's d | -2.55273 |
| 10 | Student's t | -5.70321[a] | 12.0 | <.001 | Cohen's d | -3.04849 |
| 11 | Student's t | -5.84599[a] | 12.0 | <.001 | Cohen's d | -3.12481 |
| DayAfter_Baseline | Student's t | -4.91790[a] | 12.0 | <.001 | Cohen's d | -2.62873 |
| Half_1 | Student's t | -4.20688[a] | 12.0 | 0.001 | Cohen's d | -2.24867 |
| Half_2 | Student's t | -5.59988[a] | 12.0 | <.001 | Cohen's d | -2.99326 |
| REAC_MAX | Student's t | -5.05310[a] | 12.0 | <.001 | Cohen's d | -2.70100 |
| REAC_MAX_Minusbaseline | Student's t | -3.73720[a] | 12.0 | 0.003 | Cohen's d | -1.99762 |
| REAC_% | Student's t | 1.33754 | 12.0 | 0.206 | Cohen's d | 0.71494 |
| CORT_ACCUM | Student's t | -5.14354[a] | 12.0 | <.001 | Cohen's d | -2.74934 |
| RECOVERY | Student's t | 0.00227 | 12.0 | 0.998 | Cohen's d | 0.00121 |
| RECOV_% | Student's t | 2.19864 | 12.0 | 0.048 | Cohen's d | 1.17522 |
| AUC | Student's t | -5.55036[a] | 12.0 | <.001 | Cohen's d | -2.96679 |
| AUC_BL | Student's t | -4.79259 | 12.0 | <.001 | Cohen's d | -2.56175 |
| CCL4 | Student's t | 0.96423 | 12.0 | 0.354 | Cohen's d | 0.51540 |
| TNFa | Student's t | 0.38692 | 12.0 | 0.706 | Cohen's d | 0.20682 |
| IP-10 | Student's t | 0.69571 | 12.0 | 0.500 | Cohen's d | 0.37187 |
| CCL4_Change | Student's t | -1.80433 | 12.0 | 0.096 | Cohen's d | -0.96445 |
| IP10_Change | Student's t | -1.20072 | 12.0 | 0.253 | Cohen's d | -0.64181 |
| Rest | Student's t | 1.17125 | 12.0 | 0.264 | Cohen's d | 0.62606 |
| Move | Student's t | -1.39958[a] | 12.0 | 0.187 | Cohen's d | -0.74811 |
| Eat | Student's t | 0.51944 | 12.0 | 0.613 | Cohen's d | 0.27765 |
| Jump | Student's t | -1.00614[a] | 12.0 | 0.334 | Cohen's d | -0.53781 |
| Hang | Student's t | -0.73063[a] | 12.0 | 0.479 | Cohen's d | -0.39054 |
| Sit w/o | Student's t | -0.50711 | 12.0 | 0.621 | Cohen's d | -0.27106 |
| Sit w/a | Student's t | 0.99158 | 12.0 | 0.341 | Cohen's d | 0.53002 |
| 1 | Student's t | -1.15470 | 12.0 | 0.271 | Cohen's d | -0.61721 |
| 2 | Student's t | 3.36576 | 12.0 | 0.006 | Cohen's d | 1.79908 |
| 3 | Student's t | -1.49785 | 12.0 | 0.160 | Cohen's d | -0.80063 |
| 4 | Student's t | -0.26746 | 12.0 | 0.794 | Cohen's d | -0.14296 |
| Tr | Student's t | -2.69347[a] | 12.0 | 0.020 | Cohen's d | -1.43972 |
| Tensed | Student's t | 0.17695 | 11.0 | 0.863 | Cohen's d | 0.09845 |
| Relaxed | Student's t | 0.05570 | 11.0 | 0.957 | Cohen's d | 0.03099 |
| OV | Student's t | -0.69319[a] | 11.0 | 0.503 | Cohen's d | -0.38566 |
| Far | Student's t | 1.86646 | 12.0 | 0.087 | Cohen's d | 0.99766 |
| Near | Student's t | -0.33464 | 12.0 | 0.744 | Cohen's d | -0.17887 |

Independent Samples T-Test by HPA Reactor

| | | Statistic | df | p | | Effect Size |
|---|---|---|---|---|---|---|
| Contact | Student's t | -2.57119 | 12.0 | 0.024 | Cohen's d | -1.37436 |
| Approach | Student's t | -0.60418 | 12.0 | 0.557 | Cohen's d | -0.32295 |
| Approach latency (s) | Student's t | -0.29214 | 12.0 | 0.775 | Cohen's d | -0.15616 |
| Leave | Student's t | -3.14803[a] | 12.0 | 0.008 | Cohen's d | -1.68269 |
| Leave Latency (s) | Student's t | 2.84542 | 12.0 | 0.015 | Cohen's d | 1.52094 |
| A to L Ratio | Student's t | 1.13382 | 12.0 | 0.279 | Cohen's d | 0.60605 |
| Receive Groom | Student's t | -2.67809[a] | 12.0 | 0.020 | Cohen's d | -1.43150 |
| Receive Groom (s) | Student's t | -2.65736[a] | 12.0 | 0.021 | Cohen's d | -1.42042 |
| Give Groom | Student's t | 1.06066[a] | 12.0 | 0.310 | Cohen's d | 0.56695 |
| Give Groom (s) | Student's t | 1.30326[a] | 12.0 | 0.217 | Cohen's d | 0.69662 |
| Mate | Student's t | 0.20316[a] | 12.0 | 0.842 | Cohen's d | 0.10859 |
| Self-Groom | Student's t | 0.17486 | 12.0 | 0.864 | Cohen's d | 0.09347 |
| SM | Student's t | -1.60023[a] | 12.0 | 0.136 | Cohen's d | -0.85536 |
| GD | Student's t | 0.69706 | 12.0 | 0.499 | Cohen's d | 0.37259 |
| Rest (2) | Student's t | -0.60049 | 12.0 | 0.559 | Cohen's d | -0.32097 |
| Move (2) | Student's t | 0.54509 | 12.0 | 0.596 | Cohen's d | 0.29137 |
| Eat (2) | Student's t | 0.58897 | 12.0 | 0.567 | Cohen's d | 0.31482 |
| Jump (2) | Student's t | 0.34092 | 12.0 | 0.739 | Cohen's d | 0.18223 |
| Sit | Student's t | 1.12989 | 12.0 | 0.281 | Cohen's d | 0.60395 |
| Stand | Student's t | -1.01470 | 12.0 | 0.330 | Cohen's d | -0.54238 |
| Tr (2) | Student's t | -1.07211 | 12.0 | 0.305 | Cohen's d | -0.57307 |
| Tensed (2) | Student's t | -0.88280 | 11.0 | 0.396 | Cohen's d | -0.49115 |
| Relaxed (2) | Student's t | 0.85150 | 11.0 | 0.413 | Cohen's d | 0.47373 |
| Head Scans | Student's t | -0.55081 | 12.0 | 0.592 | Cohen's d | -0.29442 |
| SM (2) | Student's t | 0.49605 | 12.0 | 0.629 | Cohen's d | 0.26515 |
| Self Groom | Student's t | 1.18032[a] | 12.0 | 0.261 | Cohen's d | 0.63091 |
| In Bucket | Student's t | 0.44699 | 12.0 | 0.663 | Cohen's d | 0.23893 |
| Shrill | Student's t | -1.05466 | 12.0 | 0.312 | Cohen's d | -0.56374 |
| BCS | Student's t | 0.00000 | 12.0 | 1.000 | Cohen's d | 0.00000 |
| Weight | Student's t | 0.29632 | 12.0 | 0.772 | Cohen's d | 0.15839 |
| Weight Proj St | Student's t | 0.63456 | 12.0 | 0.538 | Cohen's d | 0.33919 |
| Weight 6 months out | Student's t | -0.57935 | 12.0 | 0.573 | Cohen's d | -0.30968 |
| 6 M Weight Chg % | Student's t | -2.30005 | 12.0 | 0.040 | Cohen's d | -1.22943 |
| Weight Max % Chg Prev Yr % | Student's t | -0.59402 | 12.0 | 0.564 | Cohen's d | -0.31752 |
| A/G RATIO (CALC) | Student's t | 1.40892 | 12.0 | 0.184 | Cohen's d | 0.75310 |
| ALBUMIN | Student's t | 1.90888 | 12.0 | 0.080 | Cohen's d | 1.02034 |
| ALK PHOS | Student's t | -0.60921 | 12.0 | 0.554 | Cohen's d | -0.32564 |
| ALT / SGPT | Student's t | 0.65513 | 12.0 | 0.525 | Cohen's d | 0.35018 |
| ANION GAP (CALC) | Student's t | 0.91773 | 12.0 | 0.377 | Cohen's d | 0.49055 |
| AST / SGOT | Student's t | 1.67286[a] | 12.0 | 0.120 | Cohen's d | 0.89418 |
| BUN | Student's t | 0.62488 | 12.0 | 0.544 | Cohen's d | 0.33401 |

Independent Samples T-Test by HPA Reactor

| | | Statistic | df | p | | Effect Size |
|---|---|---|---|---|---|---|
| BUN/CREAT RATIO | Student's t | -1.62660 | 12.0 | 0.130 | Cohen's d | -0.86945 |
| CALCIUM | Student's t | 2.26158 | 12.0 | 0.043 | Cohen's d | 1.20886 |
| CARBON DIOXIDE | Student's t | -0.91304 | 12.0 | 0.379 | Cohen's d | -0.48804 |
| CHLORIDE | Student's t | -0.52810 | 12.0 | 0.607 | Cohen's d | -0.28228 |
| CHOLESTEROL | Student's t | 0.67922 | 12.0 | 0.510 | Cohen's d | 0.36306 |
| CPK | Student's t | 0.44520 | 11.0 | 0.665 | Cohen's d | 0.24769 |
| CREATININE | Student's t | 1.35213 | 12.0 | 0.201 | Cohen's d | 0.72274 |
| GGT | Student's t | 1.90354[a] | 11.0 | 0.083 | Cohen's d | 1.05903 |
| GLOBULIN (CALC) | Student's t | 0.06047 | 12.0 | 0.953 | Cohen's d | 0.03232 |
| GLUCOSE | Student's t | 0.32864 | 12.0 | 0.748 | Cohen's d | 0.17567 |
| LDH | Student's t | 0.55058[a] | 11.0 | 0.593 | Cohen's d | 0.30631 |
| PHOSPHORUS | Student's t | 1.41043 | 12.0 | 0.184 | Cohen's d | 0.75391 |
| POTASSIUM | Student's t | -0.69283 | 12.0 | 0.502 | Cohen's d | -0.37033 |
| SODIUM | Student's t | 0.10464 | 12.0 | 0.918 | Cohen's d | 0.05593 |
| TOTAL BILIRUBIN | Student's t | 0.00000 | 12.0 | 1.000 | Cohen's d | 0.00000 |
| TOTAL PROTEIN | Student's t | 1.25280 | 12.0 | 0.234 | Cohen's d | 0.66965 |
| TRIGLYCERIDES | Student's t | 0.98829 | 11.0 | 0.344 | Cohen's d | 0.54983 |
| BASO # | Student's t | 0.61237 | 12.0 | 0.552 | Cohen's d | 0.32733 |
| BASO % | Student's t | 0.72606 | 12.0 | 0.482 | Cohen's d | 0.38810 |
| EOS # | Student's t | -1.47029 | 12.0 | 0.167 | Cohen's d | -0.78591 |
| EOS % | Student's t | -1.72378 | 12.0 | 0.110 | Cohen's d | -0.92140 |
| Hematocrit | Student's t | -1.14985[a] | 12.0 | 0.273 | Cohen's d | -0.61462 |
| HEMOGLOBIN | Student's t | -1.31629[a] | 12.0 | 0.213 | Cohen's d | -0.70359 |
| LYMPH # | Student's t | 0.93722 | 12.0 | 0.367 | Cohen's d | 0.50097 |
| LYMPH % | Student's t | 1.03712 | 12.0 | 0.320 | Cohen's d | 0.55437 |
| MCH | Student's t | -0.83805 | 12.0 | 0.418 | Cohen's d | -0.44795 |
| MCHC | Student's t | -0.36452 | 12.0 | 0.722 | Cohen's d | -0.19484 |
| MCV | Student's t | -0.50663 | 12.0 | 0.622 | Cohen's d | -0.27081 |
| MONO # | Student's t | -0.96225 | 12.0 | 0.355 | Cohen's d | -0.51434 |
| MONO % | Student's t | -1.61378 | 12.0 | 0.133 | Cohen's d | -0.86260 |
| MPV | Student's t | 0.52281 | 12.0 | 0.611 | Cohen's d | 0.27945 |
| NEUT # | Student's t | -1.14112 | 12.0 | 0.276 | Cohen's d | -0.60995 |
| NEUT % | Student's t | -0.89371 | 12.0 | 0.389 | Cohen's d | -0.47771 |
| NRBC % | Student's t | 1.22429[a] | 12.0 | 0.244 | Cohen's d | 0.65441 |
| PLATELET COUNT | Student's t | 0.22713 | 12.0 | 0.824 | Cohen's d | 0.12141 |
| RBC | Student's t | -1.10116[a] | 12.0 | 0.292 | Cohen's d | -0.58859 |
| RDW | Student's t | 0.51489 | 12.0 | 0.616 | Cohen's d | 0.27522 |
| WBC | Student's t | 0.41831 | 12.0 | 0.683 | Cohen's d | 0.22360 |
| A/G RATIO (CALC) (2) | Student's t | -0.49692 | 11.0 | 0.629 | Cohen's d | -0.27646 |
| ALBUMIN (2) | Student's t | 1.28677 | 11.0 | 0.225 | Cohen's d | 0.71589 |
| ALK PHOS (2) | Student's t | -0.41520 | 11.0 | 0.686 | Cohen's d | -0.23100 |

Independent Samples T-Test by HPA Reactor

| | | Statistic | df | p | | Effect Size |
|---|---|---|---|---|---|---|
| ALT / SGPT (2) | Student's t | -0.18415 | 11.0 | 0.857 | Cohen's d | -0.10245 |
| ANION GAP (CALC) (2) | Student's t | 0.07528 | 11.0 | 0.941 | Cohen's d | 0.04188 |
| AST / SGOT (2) | Student's t | -0.70026[a] | 11.0 | 0.498 | Cohen's d | -0.38959 |
| BUN (2) | Student's t | -0.30105 | 11.0 | 0.769 | Cohen's d | -0.16749 |
| BUN/CREAT RATIO (2) | Student's t | -1.60244 | 11.0 | 0.137 | Cohen's d | -0.89151 |
| CALCIUM (2) | Student's t | 2.24986 | 11.0 | 0.046 | Cohen's d | 1.25171 |
| CARBON DIOXIDE (2) | Student's t | 0.53243 | 11.0 | 0.605 | Cohen's d | 0.29622 |
| CHLORIDE (2) | Student's t | 0.43019 | 11.0 | 0.675 | Cohen's d | 0.23933 |
| CHOLESTEROL (2) | Student's t | -0.82192[a] | 11.0 | 0.429 | Cohen's d | -0.45727 |
| CPK (2) | Student's t | -0.15100 | 11.0 | 0.883 | Cohen's d | -0.08401 |
| CREATININE (2) | Student's t | 1.03304 | 11.0 | 0.324 | Cohen's d | 0.57473 |
| GGT (2) | Student's t | 0.36243 | 11.0 | 0.724 | Cohen's d | 0.20164 |
| GLOBULIN (CALC) (2) | Student's t | 1.24484 | 11.0 | 0.239 | Cohen's d | 0.69257 |
| GLUCOSE (2) | Student's t | -1.90050 | 11.0 | 0.084 | Cohen's d | -1.05734 |
| LDH (2) | Student's t | 0.42608 | 11.0 | 0.678 | Cohen's d | 0.23705 |
| PHOSPHORUS (2) | Student's t | 2.18152 | 11.0 | 0.052 | Cohen's d | 1.21369 |
| POTASSIUM (2) | Student's t | 1.71325 | 11.0 | 0.115 | Cohen's d | 0.95317 |
| SODIUM (2) | Student's t | 0.69775 | 11.0 | 0.500 | Cohen's d | 0.38819 |
| TOTAL BILIRUBIN (2) | Student's t | 1.12349 | 11.0 | 0.285 | Cohen's d | 0.62505 |
| TOTAL PROTEIN (2) | Student's t | 1.66853 | 11.0 | 0.123 | Cohen's d | 0.92828 |
| TRIGLYCERIDES (2) | Student's t | 0.58422 | 11.0 | 0.571 | Cohen's d | 0.32503 |
| BASO # (2) | Student's t | -0.50383 | 11.0 | 0.624 | Cohen's d | -0.28031 |
| BASO % (2) | Student's t | -0.99871 | 11.0 | 0.339 | Cohen's d | -0.55563 |
| EOS # (2) | Student's t | 0.69494 | 11.0 | 0.502 | Cohen's d | 0.38663 |
| EOS % (2) | Student's t | 0.27794 | 11.0 | 0.786 | Cohen's d | 0.15463 |
| Hematocrit (2) | Student's t | 0.27235 | 11.0 | 0.790 | Cohen's d | 0.15152 |
| HEMOGLOBIN (2) | Student's t | -0.05042 | 11.0 | 0.961 | Cohen's d | -0.02805 |
| LYMPH # (2) | Student's t | 1.75457[a] | 11.0 | 0.107 | Cohen's d | 0.97615 |
| LYMPH % (2) | Student's t | -0.22516 | 11.0 | 0.826 | Cohen's d | -0.12527 |
| MCH (2) | Student's t | -1.18334 | 11.0 | 0.262 | Cohen's d | -0.65835 |
| MCHC (2) | Student's t | -1.38661 | 11.0 | 0.193 | Cohen's d | -0.77144 |
| MCV (2) | Student's t | -0.49230 | 11.0 | 0.632 | Cohen's d | -0.27389 |
| MONO # (2) | Student's t | -1.60e−17 | 11.0 | 1.000 | Cohen's d | -8.92e−18 |
| MONO % (2) | Student's t | -1.55871 | 11.0 | 0.147 | Cohen's d | -0.86719 |
| MPV (2) | Student's t | 0.94122 | 11.0 | 0.367 | Cohen's d | 0.52365 |
| NEUT # (2) | Student's t | 1.85130 | 11.0 | 0.091 | Cohen's d | 1.02997 |
| NEUT % (2) | Student's t | 0.92415 | 11.0 | 0.375 | Cohen's d | 0.51415 |
| NRBC % (2) | Student's t | -0.56076 | 11.0 | 0.586 | Cohen's d | -0.31198 |
| PLATELET COUNT (2) | Student's t | -0.18243 | 11.0 | 0.859 | Cohen's d | -0.10149 |
| RBC (2) | Student's t | 0.34476 | 11.0 | 0.737 | Cohen's d | 0.19181 |
| RDW (2) | Student's t | 0.51327 | 11.0 | 0.618 | Cohen's d | 0.28556 |

Independent Samples T-Test by HPA Reactor

| | | Statistic | df | p | | Effect Size |
|---|---|---|---|---|---|---|
| WBC (2) | Student's t | 2.20297[a] | 11.0 | 0.050 | Cohen's d | 1.22562 |

Note. $H_a$ $\mu_{NR} \neq \mu_{Reactive}$

[a] Levene's test is significant ($p < .05$), suggesting a violation of the assumption of equal variances

**SI Table 3.** T-Tests for each measured parameter based on HPA reactor. Behavioral variables followed by "(2)" are isolation behaviors. Blood CBC/Chem parameters are averaged or parameters followed by "(2)" are change from post-pre. Positive t-statistics indicate non-reactor > reactor. For clarification on Behavioral variable names, see Table 1.
